# Nuclear SUN2 coordinates endothelial cell-matrix interactions to regulate blood vessel homeostasis and barrier function

**DOI:** 10.64898/2026.05.18.725979

**Authors:** Pauline Bougaran, Danielle B Buglak, Alexandra Neal, Mitesh Rathod, Michaelanthony Gore, Max A Hockenberry, Aryan A Amin, Natalie Tanke, Morgan Oatley, Wesley R Legant, Ziqing Liu, James E Bear, William J Polacheck, Victoria L Bautch

**Affiliations:** Dept of Biology, The University of North Carolina at Chapel Hill, Chapel Hill, North Carolina USA; Curriculum in Cell Biology and Physiology, The University of North Carolina at Chapel Hill, Chapel Hill, North Carolina USA; Lampe Joint Dept of Biomedical Engineering, University of North Carolina Chapel Hill and North Carolina State University, North Carolina USA; Dept of Pharmacology, University of North Carolina-Chapel Hill Medical School, Chapel Hill, North Carolina, USA; Dept of Cell Biology and Physiology, UNC-Chapel Hill School of Medicine, Chapel Hill, North Carolina USA; UNC Lineberger Comprehensive Cancer Center, Chapel Hill, North Carolina USA; McAllister Heart Institute, The University of North Carolina at Chapel Hill, Chapel Hill, North Carolina USA

**Author notes:** Author for correspondence: Victoria L Bautch, PhD, Department of Biology, CB No, 3280, The University of North Carolina at Chapel Hill Chapel Hill, NC 27599, USA. Current address: Ziqing Liu, PhD, Dept of Physiology and Cardiovascular Research Center, Medical College of Wisconsin, Milwaukee, WI 53226. ACKNOWLEDGEMENTS We thank members of the Bautch lab and Dr. Steve Rogers for productive discussions. We thank the UNC High-Throughput Sequencing Facility for support. We thank Dr. Pablo Ariel, Director of the UNC Microscopy Services Laboratory, for training and support. The Andor Dragonfly microscope was funded with support from National Institutes of Health grant S10OD030223. We also thank Dr. Dan Buster for the software used in microtubule tip tracking experiments.

**Keywords:** extracellular matrix, focal adhesion, nuclear mechanotransduction, LINC complex, SUN2, microtubules

## Abstract

Vascular endothelial cells respond to environmental forces to remodel vessels during development and to achieve homeostasis, and mis-regulated responses lead to vascular dysfunction and disease. The nucleus participates in force transduction to cell-matrix junctions via the Linker of Nucleoskeleton and Cytoskeleton (LINC) complex that resides in the nuclear envelope, but how these forces are regulated and relayed is incompletely understood. We found that the LINC complex protein SUN2 is required for proper endothelial cell-matrix interactions that occur far from the nucleus and affect angiogenic expansion, vascular responses to flow, and barrier integrity. Endothelial cells lacking SUN2 had inappropriate flow responses and reduced expression of flow-mediated transcription factors *in vitro* and *in vivo*. Expression of several matrix and adhesion genes was reduced in SUN2-depleted cells, leading to defective extracellular matrix, dysmorphic focal adhesions resistant to dynamic turnover, and disturbed cell-matrix force distribution. Mechanistically, nuclear SUN2 affected dynamic regulation of the microtubule cytoskeleton that correlated with matrix metalloprotease-dependent barrier dysfunction. These findings indicate that nuclear SUN2 establishes and maintains blood vessel homeostasis by controlling microtubule-mediated effects on focal adhesion turnover and extracellular matrix properties, with implications for cardiovascular aging and diseases such as Marfan syndrome that affect vessel wall integrity.

## INTRODUCTION

Endothelial cells line the inner surface of all blood vessels and continuously experience mechanical forces from their environment. Located at the interface between circulating blood and the vessel wall, they are exposed to blood flow, including shear stress and cyclic strain [1–3], and they connect to the underlying extracellular matrix (ECM) and neighboring cells [4]. The ability of endothelial cells to detect and respond to environmental forces is essential for vascular development and homeostasis, and dysfunctional responses lead to abnormal vascular remodeling, barrier permeability, and contribute to vascular diseases such as atherosclerosis and aortic aneurysm [5].

Mechanical forces are detected and converted into biochemical signals by mechanotransducers, including cell-cell junction complexes and integrins [6–8]. When activated, these transducers initiate intracellular signaling pathways and cytoskeletal remodeling, ultimately leading to changes in cell shape, polarization, and gene expression that are relatively well-characterized [9].

The nucleus also functions as a force sensor and transducer [10–12]. The endothelial cell nucleus often protrudes into the vessel lumen, where it is subjected to mechanical drag forces from blood flow [13]. These shear forces rearrange the microtubule and actin networks and displace the nucleus relative to the MTOC (microtubule-organizing center), leading to endothelial cell polarization in response to flow [14, 15]. A crucial role for the nucleus in mechanotransduction and force generation was revealed by nuclear isolation and enucleation studies [16, 17]. The nucleus also receives mechanical stimuli from the ECM via focal adhesion linkages to the cytoskeleton [18]. The importance of nuclear mechanics in vascular function is highlighted in Hutchinson-Gilford progeria syndrome (HGPS), where premature aging due to mutation of a nuclear lamin gene results in early death from vascular dysfunction and cardiovascular complications [19]. However, the mechanisms by which mechanical cues from the microenvironment are transmitted to and from the nucleus, and how they integrate into endothelial cell responses, are not well understood.

The LINC (Linker of Nucleoskeleton and Cytoskeleton) complex physically connects the nucleus to the cytoskeleton in eukaryotic cells and mediates mechanical force transmission. Located in the nuclear envelope, the LINC complex is composed of SUN (Sad1 and UNC-84) proteins anchored in the inner nuclear membrane, and they interact with large KASH (Klarsicht, ANC-1, Syne-1 Homology, also called nesprins) proteins that span the outer nuclear membrane. On the nucleoplasmic side, SUN proteins bind the nuclear lamina, while on the cytoplasmic side they bind nesprins that connect to major cytoskeletal components, including microtubules, actin, and intermediate filaments, thereby bridging the nucleoplasm and the cytoskeleton. Most somatic cells express two SUN proteins, SUN1 and SUN2, and several nesprins [20, 21]. Nesprin-1 and-2 connect to actin via an N-terminal calponin homology domain, while the microtubule connection is mediated by motor proteins that bind nesprins near the outer nuclear membrane on the cytoplasmic side [21, 22]. SUN proteins form homo-trimers that bind both nesprin-1 and nesprin-2 to form LINC complexes. Structural and biochemical studies show that both SUN1 and SUN2 form trimeric complexes with nesprins, with SUN1 being more efficiently incorporated into LINC complexes [23–25]. Both SUN and nesprin proteins mediate force transmission across the nuclear envelope. Loss of both SUN proteins blunts traction forces, and applying direct force to nesprin-1 of isolated nuclei affects stiffness in a SUN-dependent manner [16, 17].

However, it remains unclear how different SUN complexes are assembled or utilized, and removal or over-expression of individual components often results in complex cellular changes that can be cell-type dependent [26].

LINC complexes function in vascular endothelial cells, yet how endothelial LINC-mediated regulation integrates with other environmental cues to regulate cytoplasmic structures such as cell-cell and cell-matrix junctions is not well understood. Nesprin-1-deficient HUVEC (Human Umbilical Vein Endothelial Cells) align abnormally in response to cyclic strain [27], and co-depletion of nesprin-1 and-2 decreases flow-induced tight junction protein expression [28]. Moreover, SUN proteins are important in the disease manifestation of HGPS. Cells expressing mutant lamin or HGPS patient-derived cells accumulate excess SUN1 and SUN2 at nuclear and peri-nuclear membranes, leading to increased cellular senescence, and lifespan is increased in animals expressing mutant lamin but lacking SUN1, indicating that SUN1 mediates effects of mutant lamin dysfunction in HGPS [29–31].

We previously manipulated SUN1 in vascular endothelial cells and found that it is required for proper endothelial barrier function *in vitro* and *in vivo* [32]. This work revealed that SUN1/nesprin-1 LINC complexes regulate endothelial microtubule (MT) dynamics, GEF-H1-dependent RhoA signaling, and cell-cell junction integrity. Here, we focused on understanding how SUN2 functions in endothelial cells and blood vessels, and we found that SUN2 is essential for appropriate endothelial responses to environmental cues both *in vitro* and *in vivo* in ways distinct from SUN1. SUN2 uniquely affects endothelial responses to laminar flow and regulates cell-matrix interactions far from the nucleus via focal adhesion dynamics and expression of selected ECM genes. SUN2 regulation of endothelial MT dynamics is opposite to that of SUN1, indicating that SUN2 LINC complexes normally balance SUN1 LINC complexes to regulate vascular endothelial cell function, and suggesting that this balance is compromised in cardiovascular disease.

## MATERIALS AND METHODS

### Mice

Mouse (*Mus* musculus) experiments performed in this study were approved by the University of North Carolina at Chapel Hill Institutional Animal Care and Use Committee (IACUC). *Sun2^tm1a^* mice (Strain name: C57BL/6N-Atm1Brd Sun2*^tm1a(EUCOMM)Hmgu^*/BayMmucd, MMRRC ID: #37788) were obtained from the Mutant Mouse Resource & Research Centers (MMRRC). FlpO-B6N-Albino (Rosa26-FlpO/+) mice were obtained from the UNC Animal Model Core. *Cdh5^CreERT2^* mice were generated by Dr. Ralf Adams and obtained from the Cancer Research UK [33]. To obtain the *Sun2* floxed allele (*Sun2^fl^*), *Sun2^tm1a^* mice were bred with Flp-recombinase-expressing mice (FlpO-B6N-Albino (Rosa26-FlpO/+)). The *Sun2^tm1a^* allele was detected by genomic PCR using primers amplifying the LacZ insertion (see Table S1 for primer details). Flp-mediated recombination removed the *LacZ* cassette and generated the *Sun2^fl^*allele, in which exons 3 and 4 of *Sun2* are flanked by *loxP* sites (**Supp. Fig. 1G**). To generate an endothelial cell-specific conditional knockout, *Sun2^fl/fl^*mice were crossed with *Cdh5-Cre^ERT2^* mice to generate *Sun2^fl/fl^*;*Cdh5Cre^ERT2^*pups. Cre recombination was induced by intraperitoneal injection of pups at postnatal days (P) P1, P2 and P3 with 50µL of 1mg/mL tamoxifen (Sigma-Aldrich, #T5648) dissolved in sunflower seed oil (Sigma-Aldrich, #S5007). For genotyping, tail clip or toe tissue samples were incubated in alkaline lysis buffer (25mM NaOH; 0.2mM Disodium EDTA) at 92°C for 1h, then an equal volume of neutralizing solution (40mM Tris-HCl) was added. Genomic PCR was performed to confirm *Sun2* allele excision and *Cdh5-Cre^ERT2^*expression. Primer sequences are listed in **Table S1**.

### En face aorta staining

Tamoxifen-injected pups were euthanized at P7 and perfused through the left ventricle with 3mL of 4% paraformaldehyde (PFA, Electron Microscopy Sciences, #15713).

Aortas were collected, post-fixed in 4% PFA for 30min, washed in 1x PBS, carefully cleaned of surrounding tissue, and opened longitudinally. Aortas were permeabilized/blocked for 1h at room temperature (RT) with gentle rocking in blocking buffer (10% donkey serum (Sigma-Aldrich, #D9663), 0.5% Triton X-100 (Sigma-Aldrich, #T8787), and 0.01% sodium deoxycholate (Sigma-Aldrich, #D6750)) in 1X PBS.

Tissues were incubated overnight at 4°C with primary antibodies diluted in a 1:1 solution of blocking buffer and 1x PBS, with rocking. After 3X washes with 1x PBS, aortas were incubated for 1h at RT with secondary antibodies and DAPI (Roche, #10236276001) diluted in 1:1 blocking buffer:PBS solution. After 3X washes with 1x PBS, samples were flat-mounted using Prolong Diamond Antifade Mounting Media (ThermoFisher Scientific, #P36961). The primary and secondary antibodies used for aorta staining are listed in **Table S2**.

### Retina staining

Tamoxifen-injected pups were euthanized at P7, and eyes were collected, fixed in 4% PFA for 1h at RT, and washed in 1x PBS. Prior to staining, retinas were dissected and permeabilized/blocked for 1h at RT, with gentle rocking in blocking buffer, as described for *en face* aorta staining. Tissues were then incubated overnight at 4°C with Isolectin-B4 Alexa Fluor 488 (1:100, I21411, Thermo Fisher) diluted in a 1:1 solution of blocking buffer and 1x PBS, with gentle rocking. After 3 washes with 1x PBS, retinas were flat-mounted using Prolong Diamond Antifade Mounting Media.

### Zebrafish

Zebrafish (*Danio rerio*) experiments were approved by the University of North Carolina at Chapel Hill Institutional Animal Care and Use Committee (IACUC). *Tg:(fli:LifeAct-GFP)* zebrafish line was a gift from Dr. Wiebke Herzog. The *sun2*^sa16555^ fish were obtained from the Zebrafish International Resource Center (ZIRC). For genotyping, fin clip (adult fish) or tissue (euthanasized embryos) was incubated in alkaline lysis buffer at 95°C for 1h, then an equal volume of neutralizing solution was added. Genomic PCR was performed using primers to amplify the target region of the *sun2* gene using the following primers (see **Table S1** for primer details). PCR products were purified using the GeneJet PCR Purification Kit (ThermoFisher Scientific, #K0702), per the manufacturers’ protocol, and purified DNA was Sanger sequenced (GENEWIZ) using the forward primer.

Morphant fish were generated by injecting 2.5-5 ng of non-targeting (NT) morpholino or *sun2* morpholino into the yolk of *Tg:(fli:LifeAct-GFP)* embryos at the one-cell stage (see **Table S3** for morpholino sequences). Morpholinos were diluted to desired concentration in nuclease-free water (Invitrogen, #AM9937) containing 0.04% phenol red (Sigma-Aldrich, #P0290) for visualization during injection. Morphant or mutant embryos were maintained in E3 medium at 28.5°C until 36hpf, then dechorionated, fixed overnight in 4% PFA at 4°C, rinsed 3X with 1x PBS, mounted using Prolong Diamond Antifade mounting media, and sealed with petroleum jelly.

### Cell culture, siRNA and plasmid transfection

Human Umbilical Vein Endothelial Cells (HUVEC, Lonza, #C2519A) were cultured at 37°C in Endothelial Basal Medium-2 (EBM-2, Lonza, #CC3156) supplemented with Endothelial Cell Growth Medium-2 BulletKit (EGM2, Lonza, #CC4176) and 1X antibiotic-antimycotic (Gibco, #15240062). Cells at passage 3-7 were used for experiments.

*siRNA experiments*: HUVEC were incubated with non-targeting (NT) (Dharmacon, #D-001810-10-20), SYNE1 (Dharmacon, #M-014039-02-0005), or SUN2 siRNAs (Life technologies, #4392420, s24467) (**Table S3**) using Lipofectamine 3000 (ThermoFisher Scientific, #L3000015) according to manufacturers’ protocol, at a final concentration of 40nM, for 18-24h at 37°C, then incubated in fresh medium for 24h. For immunofluorescence experiments (except for flow conditions), cells were seeded onto glass-bottom slides (Ibidi, #80807) coated with 5µg/mL fibronectin (Sigma-Aldrich, #F2006).

*Plasmid transfection for live imaging:* HUVEC were transfected with plasmid constructs using the Amaxa Nucleofector^®^ II system. Confluent monolayers were trypsinized with 0.05% Trypsin-EDTA (Gibco, #25300-054) for 3 min at 37°C, collected in fresh complete medium, pelleted at 1200 rpm for 5 min, then resuspended in 100µL of Nucleofector solution (Lonza, #VBP-1002). 2.5µg of plasmid was added to the cell suspension, and electroporation was performed (D-005 program). Cells were immediately transferred into fresh EGM-2 medium and seeded onto fibronectin-coated glass-bottom slides (Ibidi, #80807). Media was replaced 4h post-electroporation and live imaging was performed 48 hr later. For microtubule tip-tracking, EB3-mNeonGreen construct was a gift from Daniel Gerlich (Addgene plasmid #191761; http://n2t.net/addgene:191761; RRID: Addgene_191761) [34]. For focal adhesion dynamic analysis, the following constructs were used: pRK GFP Paxillin (gift Kenneth Yamada (Addgene plasmid #50529; http://n2t.net/addgene:50529; RRID: Addgene_50529) and Vinculin-venus (gift Martin Schwartz (Addgene plasmid #27300; http://n2t.net/addgene:27300; RRID: Addgene_27300) [35].

### Flow experiments

All laminar flow experiments were performed using the ibidi pump system at 15dyn/cm^2^ for 72h, unless otherwise stated.

*Ibidi pump system*: Flow experiments were performed using an Ibidi pump system (Ibidi, #10902) as described [36]. HUVEC were seeded onto Ibidi flow chambers (Ibidi, µ-Slide I Luer I 0.4 mm, #80176) pre-coated with 5µg/mL fibronectin at a density of 225,000 cells/slide, in EBM-2 medium supplemented with 2% fetal bovine serum (FBS, Gibco, #16-140-089), 1% antibiotic-antimycotic, and 1% Nystatin (Sigma-Aldrich, #N1638).

After 4h, laminar flow (15 dyn/cm^2^, 72h) was initiated. Static (non-flow) control slides were maintained under identical medium conditions for 48h before fixation or lysis for RNA extraction.

*Orbital shaker:* For Western Blot, EdU labeling, and drug addition, laminar flow was applied using an orbital shaker (VWR, Standard Analog Shaker), as described [36]. HUVEC were seeded in 6-well plates at a density of 5 x 10*^5^* cells/well and cultured for 24h before exposure to shear stress for 24h. To ensure consistent sampling, cells located at the center of each well were scraped and removed using the same circular template for all experiments, and only peripheral cells exposed to uniaxial flow were collected for further analysis.

### Microfluidic devices

*3D Microvessel Formation:* Microvessel formation was carried out following previously described procedures [37]. To produce microvessels, confluent HUVEC were treated with 0.05% (w/v) trypsin-EDTA (Thermo Fisher), then resuspended in EGM-2 to achieve a final concentration of 1.8×10^6^ cells/mL. Before seeding into microfluidic devices, gel-filling pipette tips were inserted into the hydrogel ports and gently aspirated 3 times to remove residual media and hydrogel, creating a pressure gradient across the vessel wall. Media was removed from the inlet and outlet ports, then 60μL of the cell suspension was introduced into one port, and 50μL into the other to generate a slight pressure differential, promoting cell flow into the microchannel. The devices were inverted one per minute for 8 min, then placed upside down in a humidified incubator for 6–8 min, followed by re-inversion and a second incubation for 6–8 min. Devices were then transferred to a rocker and oscillated at 5 cycles/min for 2–4 hr before replacing media with fresh EGM-2. Devices remained on the rocker overnight and were visually inspected the next day. If the monolayer did not fully cover the vessel, the seeding was repeated.

*3D Microvessel Permeability Assay*: The diffusive permeability of microvessels was determined as previously described [38]. Briefly, EGM-2 was supplemented with 10 μg/mL of 70 kDa Texas Red Dextran (Sigma-Aldrich) and introduced into one media port. Images of the vessel’s mid-plane were taken using a laser scanning confocal microscope (Olympus FV3000) with a 10x, 0.4 NA objective every 10 sec over a period of 2 min. Dextran extravasation was evaluated at the final time point. Vessel regions were first manually defined, and the area fraction of the thresholded dextran signal outside these regions was measured using Fiji.

### Traction force microscopy

*Generation of compliant PDMS substrates:* Compliant PDMS substrates for traction force microscopy (TFM) were generated as previously described [39]. Briefly, Sylgard™ 527 (Dow #1696742) was mixed thoroughly at a 1:1 ratio (Part A:Part B by weight), yielding substrates with an approximate Young’s modulus of 2 kPa, and degassed for 15 min in a tabletop vacuum chamber. 50 µL of the mixture was deposited onto a plasma-cleaned #1.5, 25 mm glass coverslip and spin-coated (Laurell WS-650-23) for 1 min at 2000 rpm to produce substrates ∼50 μm thick. Substrates were cured overnight at 60 °C, then transferred to a cell culture plastic 6-well plate and incubated in 10% (v/v) APTES (ThermoFisher Scientific, #430941000) in ethanol with shaking on a benchtop shaker for 30 min. After washing 3X with 100% ethanol, a solution containing 1:1000 carboxylated polystyrene fluorescent beads (ThermoFisher Scientific, #F8801) and 1 mg/mL EDC (ThermoFisher Scientific, #E7750) in deionized water was added and incubated for 1 h with rocking. After washing 3X with deionized water, substrates were coated with fibronectin and sterilized by placing 1 mL of 10 μg/mL fibronectin (Gibco, #33010018) in a fresh sterile 6-well plate, inverting the substrates onto the solution, and exposing them to UV light for 30 min in a tissue culture hood. After sterilization, substrates were inverted again and washed 3X with 1x PBS prior to cell seeding.

*Imaging of cells on substrates:* Cells were plated at a density of ∼3000 cells per substrate. Substrates were transferred to a 35-mm metal imaging dish, and culture medium containing cells was added. Dishes were covered with a sterile 35 mm lid and incubated overnight. Substrates were imaged next day on a Nikon Ti2 inverted widefield microscope equipped with a live-cell chamber maintained at 37 °C and 5% CO_₂_.

Imaging was performed using a 60× SR Plan Apo IR water immersion objective (NA 1.27). Fluorescence images were acquired using a 560 nm LED (10% power, 100 ms exposure), and brightfield images were collected (5% condenser lamp power, 100 ms exposure). After imaging, cells were removed by addition of 100 μL of 10% (w/v) SDS for 5 min, then acquiring a reference image of the relaxed bead state.

*Calculation of traction force parameters:* Cell-induced traction forces were computed as previously described [40]. TFM image sequences were converted to single-frame TIFF files and processed using μ-inferforce [41]. Subpixel drift correction was performed using Efficient Subpixel Registration. Displacement fields were calculated using high-resolution bead subsampling, with a bead selection threshold (alpha) of 0.05 and a template size of 21 pixels. Traction forces were reconstructed using Fourier Transform Traction Cytometry (FTTC) with a regularization parameter of 0.0001, assuming a substrate modulus of 2 kPa and a thickness of 50 μm. Contractile energy and force dipole ratio were calculated as described previously [42], where contractile energy is defined as the total strain energy and the force dipole ratio is the ratio of the major to minor dipole components.

### Cell treatments and functional assays

*Cell treatments:* Confluent HUVEC were treated with 10nM nocodazole (Sigma-Aldrich, #M1404, in DMSO) for 4h at 37°C. HUVEC were treated with 5µM Marimastat (Sigma-Aldrich, #M2699, in DMSO) or with 25µM Ilomastat (MedChemExpress, #HY-15768, in DMSO) for 24h at 37°C.

*EdU Labeling:* HUVEC were seeded on 6-well glass bottom plates (Cellvis, #P06-1.5H-N) pre-coated with 5µg/mL fibronectin, at a density of 5×10***^5^*** cells per well. Orbital flow was applied for 24 h and EdU diluted in fresh complete medium was added for the last hour. Cells were rinsed once in PBS and fixed with 4% PFA for 10 min at RT. EdU labeling was performed according to the Click-IT EdU 594 protocol (Invitrogen, #C20337) and subsequent immunostaining was performed as described above.

*Decellularization:* HUVEC were seeded at 1.5×10^4^ cells/well in glass-bottom slides (Ibidi, #80807) pre-coated with 5µg/mL fibronectin. Five days after seeding, confluent monolayers were rinsed 1x with PBS and treated with decellularization solution (0.5% Triton X-100, 20mM NH_4_OH (Sigma-Aldrich, #338818)) diluted in PBS for 3 min at 37°C. Sample were rinsed incubated 3X with PBS and with 100U/mL DNAse I (Affymetrix/USB, #14340) in PBS for 45 min at 37°C. Subsequently, fresh siRNA-treated HUVEC were seeded onto the cell-derived ECM at 1×10^5^ cells/well and fixed 4h post-seeding.

*Biotin matrix labeling assay*: Biotinylated matrix labeling was adapted from Dubrovski et al. [43] and performed as previously described [32, 44]. Briefly, fibronectin (0.1mg/mL) was biotinylated by incubation with 0.5mM EZ-Link Sulfo-NHS-LC-Biotin (ThermoFisher Scientific, #A39257) for 30 min at RT. Glass-bottom slides (Ibidi, #80807) were then coated with 5µg/mL biotinylated fibronectin and seeded with HUVEC. Upon reaching confluence, monolayers were incubated with 25µg/mL Streptavidin-488 (Invitrogen, #S11223) for 3 min at 37°C, then fixed with 4% PFA for subsequent immunofluorescence analyses.

*Real time cell analysis:* Cell adhesion and monolayer integrity were assessed using the xCELLigence Real-Time Cell Analyzer (RTCA, Acea Biosciences/Roche Applied Science). HUVEC were seeded at 3X10^5^ cells/well onto microelectrodes of the E-plate 16 PET (Agilent, #300600890). Impedance-based cell index was recorded every 2 min for 24h.

### Immunofluorescence

HUVEC were rinsed with PBS and fixed with ice-cold methanol for 10min at RT to visualize microtubules. For all other experiments, cells were rinsed with PBS, fixed with 4% PFA for 10 min at RT, and permeabilized with 0.1% Triton X-100 (diluted in PBS) for 10 min at RT. Cells were then incubated for 1hr at RT with rocking in blocking buffer (5% FBS, 2% antibiotic-antimycotic, and 0.1% sodium azide (Sigma-Aldrich, #S2002)) diluted in PBS then incubated overnight at 4°C with primary antibodies diluted in blocking buffer with rocking. After 3X washes with PBS (5min each), cells were incubated for 1hr at RT with secondary antibodies and DAPI diluted in blocking buffer.

After 3X washes with PBS (5min each), slides were stored at 4°C until imaging. Primary and secondary antibodies used for HUVEC immunofluorescence are listed in **Table S2**.

### Western Blotting

HUVEC were scraped into RIPA buffer composed of 0.1% SDS (Fisher scientific, #BP166-500), 0.5% sodium deoxycholate (Sigma-Aldrich, #D6750), and 1% IGEPAL® (Sigma-Aldrich, #I8896) supplemented with 1x protease/phosphatase cocktail (Cell signaling, #5872S), and centrifuged at 1×10^4^g at 4°C for 10 min. Total protein was determined using the Bio-Rad Protein Assay Dye Reagent Concentrate (Bio-Rad, #5000006). Lysates were reduced in sample loading buffer with 0.1M DTT (dithiothreitol (ThermoFisher Scientific, #R0861) and boiled for 5 min at 100°C. Reduced samples (10µg) were separated on a 10% stain-free polyacrylamide gel (Bio-Rad, #1610183) and transferred to a PVDF membrane (Bio-Rad, #1620177) on ice. Membranes were blocked in OneBlock™ blocking buffer (Genesee Scientific, #20-313) for 1 hr at RT and incubated with primary antibodies diluted in OneBlock™, overnight at 4°C with rocking. After 3X washes (5min each) with PBS-Tween (0.1% Tween-20 (Sigma-Aldrich, #P9416) in PBS), membranes were incubated with secondary antibodies diluted in OneBlock™ for 1h at RT with rocking. After 3X washes with PBS-Tween (5min each), Immobilon Forte Western HRP Substrate (Sigma-Aldrich, #WBLUF100) was added for 30 sec to 2 min and blots were exposed for 8 sec. Primary and secondary antibodies used for Western Blot analysis are listed in **Table S2**.

### RT-qPCR

Confluent HUVEC were rinsed with PBS, collected in TRIzol™ (Invitrogen, #15596018), and total RNA was isolated according to manufacturers’ instructions. RNA concentration and purity were determined using a NanoDrop spectrophotometer (ThermoFisher Scientific, NanoDrop One). Complementary DNA (cDNA) was synthetized from 500ng of total RNA using the iScript™ cDNA Synthesis kit (Bio-Rad, #1708891) and diluted 1:5 in nuclease-free water. RT-qPCR was performed using iTaq™ Universal SYBR^®^ Green SuperMix (Bio-Rad, #1725121) on an Applied Biosystems QuantStudio 6 Flex Real-Time PCR System. Relative expression levels were calculated using the 2^(Ct of gene – Ct of GAPDH)^ and fold changes relative to control samples were determined using the ΔΔCt method. Primer sequences used for RT-qPCR analysis are provided in **Table S1**.

### Bulk RNA Sequence Analysis

HUVEC were exposed to laminar flow (15d/cm^2^, 72h). Two slides/condition (static or flow) were combined in TRIzol™ and 3 biological replicates collected. Stranded libraries were prepared using KAPA mRNA HyperPrep Kit (Roche. #7961901001), sequenced using NovaSeq S1 at the UNC Sequencing Core and data was analyzed as described previously [32, 36]. 2–3×10^7^ 50 bp paired-end reads per sample were obtained and mapped to human genome GRCh38 with STAR using default settings [45]. Mapping rate was over 80% for all samples, and gene expression was determined with Htseq-count using the union mode [46]. Differential expression analysis was performed with DESeq2 [47] using default settings in R, and lists of differentially expressed genes were obtained (p adjusted <0.1). Gene ontology analysis was performed using enrichGO function in the R package clusterProfiler.

### Image Acquisition and Analysis

Unless otherwise specified, imaging was performed using an Olympus confocal laser scanning microscope (Fluoview FV3000, IX83) with 405, 488, 561, 640 nm lasers, and a UPlanSApo 40x silicone-immersion objective (NA 1.25). Imaging of the mouse retina was performed using a UPlanSApo 10x air objective (NA 0.40). Images were acquired at RT for fixed samples, using the Olympus Fluoview FV31S-SW software with identical acquisition settings applied across all conditions. All image analysis was performed using Fiji software [48].

*Cell axis ratio:* Confocal z-stacks of VE-cadherin-labeled HUVEC or mouse aortas were processed in Fiji to generate maximum intensity projections. Cell outlines were manually drawn based on junctional staining **(Supp. Fig. 1C)**, and the major-to-minor axis ratio was calculated using the “Fit ellipse” and “Shape descriptors” functions.

*Junction linearity:* Linearity was measured manually in Fiji, by measuring the total length of a junction with the segmented line tool and using the straight-line tool to determine the Euclidean distance between the endpoints **(Supp. Fig. 1H)**. The ratio of Euclidean distance to the total junction length was calculated.

*Elastin breaks:* Breaks were quantified as previously described [49] using the autofluorescence signal of elastin fibers. The number of breaks in the first elastin sheet, located between the endothelial cells and the first vascular smooth muscle cell layer, was quantified manually in Fiji.

*Cell polarity:* Maximum intensity projections of GM130 (Golgi) and DAPI (nucleus) staining were generated using Fiji. The polarity vector was defined as the line connecting the center of the nucleus to the center of the Golgi, and the angle between this vector and the flow direction was measured manually using the Angle tool in Fiji (**Supp. Fig. 1F**). Cells with an angle between 0 and 60° were classified as polarized with the flow, those between 60° and 120° as side and those between 120° and 180° as polarized against the flow direction.

*EdU quantification:* Confocal images of EdU-labeled HUVEC were processed in Fiji, and maximum intensity projections were generated. EdU^+^ nuclei were counted manually, and a mask of DAPI-stained nuclei was created to determine total nuclei/field. The percentage of EdU^+^ cells was calculated as the ratio of EdU^+^ nuclei to total nuclei.

*KLF4 and SUN2 nuclear intensity quantification in the descending aorta:* Confocal z-stacks of stained mouse aortas were processed in Fiji to generate maximum intensity projections of the endothelium and to exclude the underlying smooth muscle cells. The DAPI channel was then used as a nuclear mask, and nuclear fluorescence intensity was measured for each cell, excluding cells at the edges.

*Retinal vessel analysis:* The radial expansion of the retinal vasculature was determined by dividing the distance from the optic nerve to the vascular front by the distance from the optic nerve to the tissue edge, with four measurements averaged per retina.

Vascular density was quantified by dividing the vascular area, determined by applying a threshold to the IB4 staining using Fiji, by the total retina area.

*Microtubule tip-tracking*: HUVECs expressing the EB3-mNeonGreen construct were imaged using an Andor Dragonfly spinning-disk confocal microscope equipped with a 37°C humidified CO_2_ incubator. Images were acquired with a Zyla Plus 4.2 MP sCMOS camera using a HC PL APO 63X/1.4 Oil objective. Time-lapse imaging was performed with 470ms exposure per frame for 2 min, capturing a total of 240 frames. Microtubules of the cell retracting edge were tracked using Manual Tracking with Trackmate plugin in Fiji, for 120 frames (60 sec). Track information was acquired, as previously described [32], from the x and y coordinates using a custom algorithm in Visual Basic in Excel provided by Dr. Dan Buster at the University of Arizona.

*Focal adhesion imaging:* HUVEC expressing Paxillin-GFP or Vinculin-venus constructs were imaged using an Olympus FV3000 equipped with a stage-top incubator (TOKAI HIT, WSKM) at 37°C and with humidified CO_2_. Images were acquired using a UPlanSApo 20× oil-immersion objective (NA 0.58) every 4 min for 40 min. To analyze focal adhesion dynamics, temporal projections of individual focal adhesions were generated using Fiji. Each focal adhesion was divided into three equal regions corresponding to the start, middle, and end. Red, blue and green channels were separated in Fiji and the mean gray value was measured for each region. The red-to-green ratio was then calculated along the focal adhesion axis, and the slope from the front to the start to the end was quantified.

### Statistics

All statistical analyses and graph generation were performed using Prism 10 software (GraphPad Software). A two-tailed unpaired Student’s t-test was used to determine statistical significance between two experimental groups, and one-way ANOVA with Tukey post hoc test was used to compare three or more groups. A Χ^2^ test was performed to evaluate differences in the distribution of cells among different polarity categories.

## RESULTS

### Nuclear SUN2 is required for proper endothelial cell responses to blood flow and vascular expansion

SUN protein-containing LINC complexes that reside at the nuclear membrane influence cellular processes and responses far from the nucleus by poorly understood mechanisms. Here, we examined how SUN2 functions in EC responses to environmental cues, focusing first on flow-mediated responses. Since disrupting LINC complexes globally causes cell detachment under laminar flow [50], but loss of SUN1 does not affect cell elongation in response to homeostatic flow [32], we hypothesized that SUN2 is necessary for EC adaptation to fluid shear stress imparted by laminar blood flow. Primary HUVEC were depleted for SUN2 via siRNA (**Supp. Fig. 1A-B**), then subjected to homeostatic laminar flow (15d/cm^2^, 72h unless otherwise stated). SUN2-depleted EC exhibited abnormally increased elongation along the flow vector compared to controls (**Fig. 1A-B; Supp. Fig. 1C**), indicating an inappropriate response of hyper-elongation. A similar hyper-elongated cell shape was observed in SUN2-depleted EC cultured under static conditions (**Supp. Fig. 1D-E**), revealing that EC depleted for SUN2 have abnormal responses to environmental cues, and that these responses include abnormal flow elongation. Moreover, SUN2-depleted EC under flow were more randomly polarized to the flow vector compared to controls, as defined by the nucleus-Golgi axis (**Fig. 1C-D, Supp. Fig. 1F**). Since laminar flow promotes EC quiescence [51], we next assessed cell proliferation. EdU incorporation is a readout of cells in the S-phase of the cell cycle, and SUN2 depletion under laminar flow significantly increased this parameter compared to controls (**Fig. 1E-F**), indicating that SUN2 is required for flow-mediated quiescence in EC. Consistent with loss of proper flow responses, SUN2 depletion also reduced flow-induced mRNA expression of two transcription factors that mediate flow responses, *KLF2* and *KLF4*, as well as KLF4 protein expression (**Fig. 1G-I**). Collectively, these findings show that nuclear SUN2 is necessary for proper EC responses to laminar flow and suggest that SUN2 also modulates endothelial responses to non-flow environmental cues.

**Figure 1.**
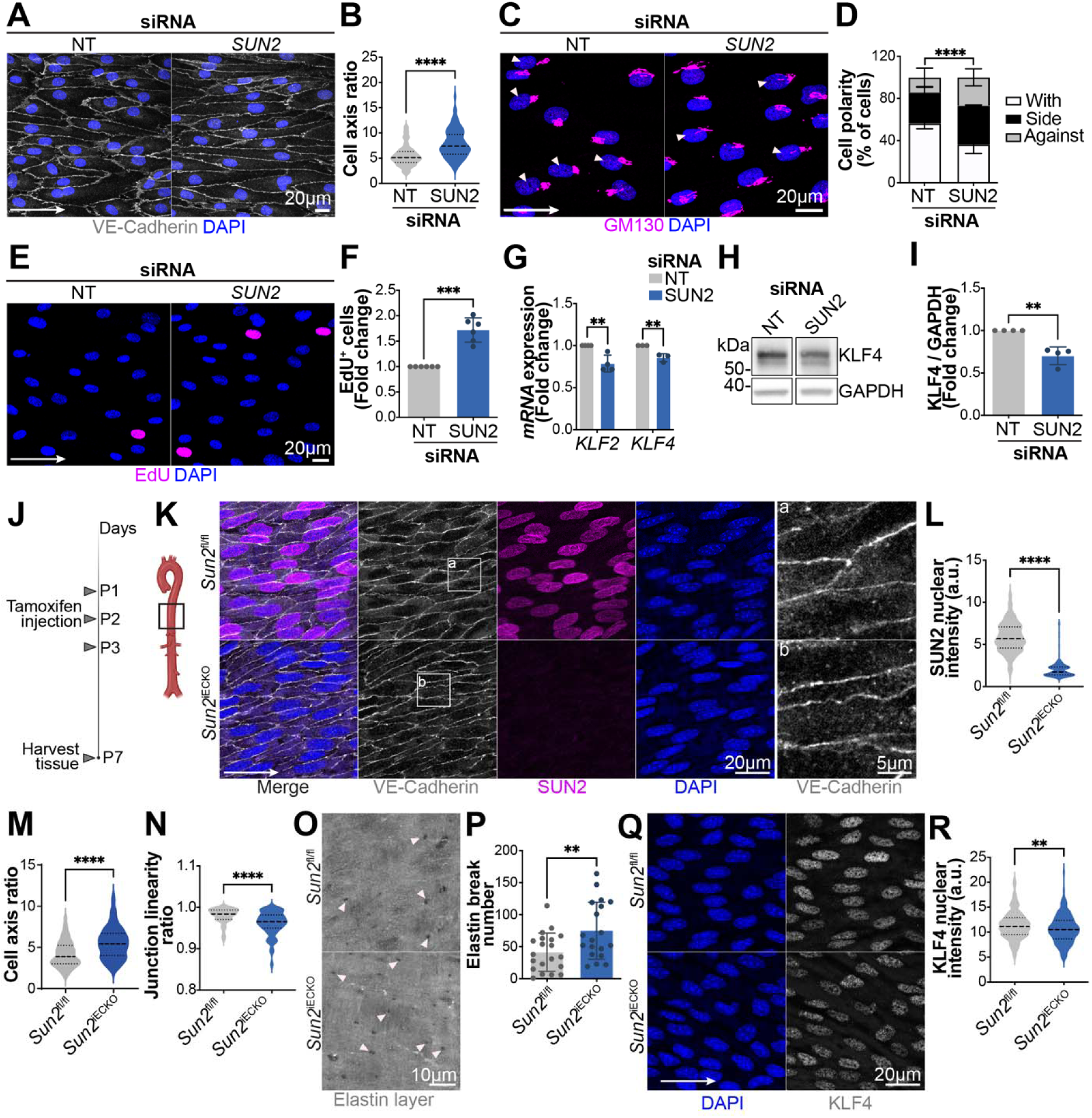
**Nuclear SUN2 regulates endothelial cell flow responses in cells and *in vivo***. **(A, C, E)** Confocal images (maximum intensity projections) of HUVEC treated with siRNA as indicated and exposed to laminar flow (**A, C**: 15 d/cm^2^, 72h; **E**: 15d/cm^2^, 24h, orbital), then fixed and stained as indicated. White triangles in C indicate cells polarized with the flow direction. White arrow, flow direction. **(B, D, F)** Quantification of indicated parameters. Scale bar, 20 µm. (n=3 biological replicates). **(G)** RT-qPCR of indicated genes in HUVEC treated as indicated and exposed to laminar flow. (n ≥ 3 experiments) **(H, I)** Western Blot and quantification of KLF4 protein levels (normalized to GAPDH) in HUVEC treated as indicated and exposed to laminar flow (15 d/cm^2^; 24h; orbital) (n=4 biological replicates). **(J)** Diagram of tamoxifen regimen. **(K, Q)** Confocal images (maximum intensity projections) of *en face* preparation of descending thoracic aorta from mice of the indicated genotypes and stained as indicated. Scale bar, 20 µm **(K: a-b)** EC junctions shown at higher magnification on far right. Scale bar: 5µm **(L-N, R)** quantification of indicated parameters from aortas of indicated genotypes (n≥ 3 aortas of each genotype). **(O)** *En face* confocal images of autofluorescent internal elastic lamina in the descending thoracic aorta of mice of indicated genotypes. White triangles indicate the elastin breaks. Scale bar, 10 µm. **(P)** Quantification of breaks in the internal elastic lamina of mice of the indicated genotypes. n=5 aortas of each genotype. Statistical analyses were performed using two-tailed Student’s t-tests, except for **(D),** which was analyzed using a χ² test. **, p ≤0.01; ***, p ≤ 0.001; ****, p ≤ 0.0001.

To assess whether SUN2 similarly influences endothelial flow responses *in vivo*, we produced a *Sun2^fl/fl^* mouse line and used it to generate a tamoxifen-inducible, endothelial-specific *Sun2* knockout mouse (*Sun2^fl/fl^*;Cdh5-Cre^ERT2^), referred to as *Sun2^iECKO^* (**Supp. Fig. 1G**). *Sun2* deletion in EC was induced by tamoxifen injection from postnatal (P) days 1-3 (**Fig. 1J**), and *en face* immunofluorescence of the P7 descending aorta confirmed endothelial loss of SUN2 protein expression in *Sun2^iECKO^* mice compared to littermate controls (*Sun2^fl/fl^*) (**Fig. 1K-L**). The endothelial cell axis ratio was significantly increased in *Sun2^iECKO^* aortas compared to controls, revealing that the inappropriate hyper-elongation response observed in HUVEC also occurs *in vivo* (**Fig. 1K, M**). Additionally, loss of *Sun2* affected endothelial cell-cell junctions: control aortas had mostly linear endothelial junctions, while *Sun2^iECKO^* mice had irregular, wavy endothelial junctions with significantly reduced linearity (**Fig. 1Ka-b, N**; **Supp. Fig. 1H**). Interestingly, a similar junctional phenotype was reported in mice carrying a mutation in the *Fbn1* (fibrillin-1, *Fbn1^C1041G/+^*) gene that models Marfan syndrome, a connective tissue disorder characterized by extracellular matrix (ECM) remodeling and degradation leading to aortic aneurysm [49]. Since Marfan patients and mice also exhibit fragmentation of elastic fibers within the aortic wall, we examined the elastin layer of the *en face* aortas and found significantly more elastin breaks in *Sun2^iECKO^* aortas compared to controls (**Fig. 1O-P**), similar to the phenotype of mice carrying the *Fbn1* mutation associated with Marfan syndrome. Moreover, nuclear KLF4 immunostaining was reduced in EC of *Sun2^iECKO^* aortas compared to control littermates (**Fig. 1Q-R**), consistent with *in vitro* data and indicating compromised EC flow responses with *Sun2* loss *in vivo*. These data collectively demonstrate that nuclear SUN2 regulates EC flow-induced responses in culture and *in vivo*. Additionally, endothelial loss of *Sun2* is associated with ECM and junctional defects that resemble those observed in Marfan syndrome and aortic aneurysm.

Since endothelial loss of *Sun2* was associated with abnormal responses to environmental cues and ECM organization, we next investigated whether nuclear SUN2 affected vascular network development and vessel morphology *in vivo*. The retinal vascular network of P7 *Sun2^iECKO^* mice had significantly reduced radial expansion compared to littermate and Cre-only expressing controls, while vascular density was not significantly affected (**Supp. Fig. 2A-C**). We next used zebrafish expressing a *Tg(fli:LifeAct-GFP)* reporter to visualize the EC actin cytoskeleton. Analysis of intersegmental vessel (ISV) development in embryos at 36 hours post-fertilization (hpf) revealed that s*un2* depletion via morpholino (MO) injection significantly impaired ISV growth and connection to the dorsal longitudinal anastomotic vessel (DLAV) (**Supp. Fig. 2D-E),** and comparable trends were observed using a *sun2*^-/-^ mutant zebrafish line (**Supp. Fig. 2F-G)**. Thus, Sun2 regulates vascular endothelial cell flow responses and vascular expansion during vascular development.

### Nuclear SUN2 regulates ECM-related gene transcription and properties

To further examine how SUN2 affects endothelial cell properties, we assessed gene expression via bulk RNA sequencing of HUVEC under flow. Expression analysis identified 116 differentially expressed genes between *SUN2-*depleted EC and controls, including 21 upregulated and 95 downregulated genes. Gene ontology (GO) analysis revealed significant enrichment in pathways related to cell-substrate adhesion and ECM organization (**Fig. 2A-B**). Although heatmap visualization of a panel of ECM-associated genes [52–54] did not reveal global dysregulation of this gene set with SUN2 depletion (**Supp. Fig. 3A)**, a subset of ECM and cell-adhesion genes downregulated in SUN2-depleted endothelial cells was identified within the GO-enriched pathways **(Fig. 2C)**.

**Figure 2.**
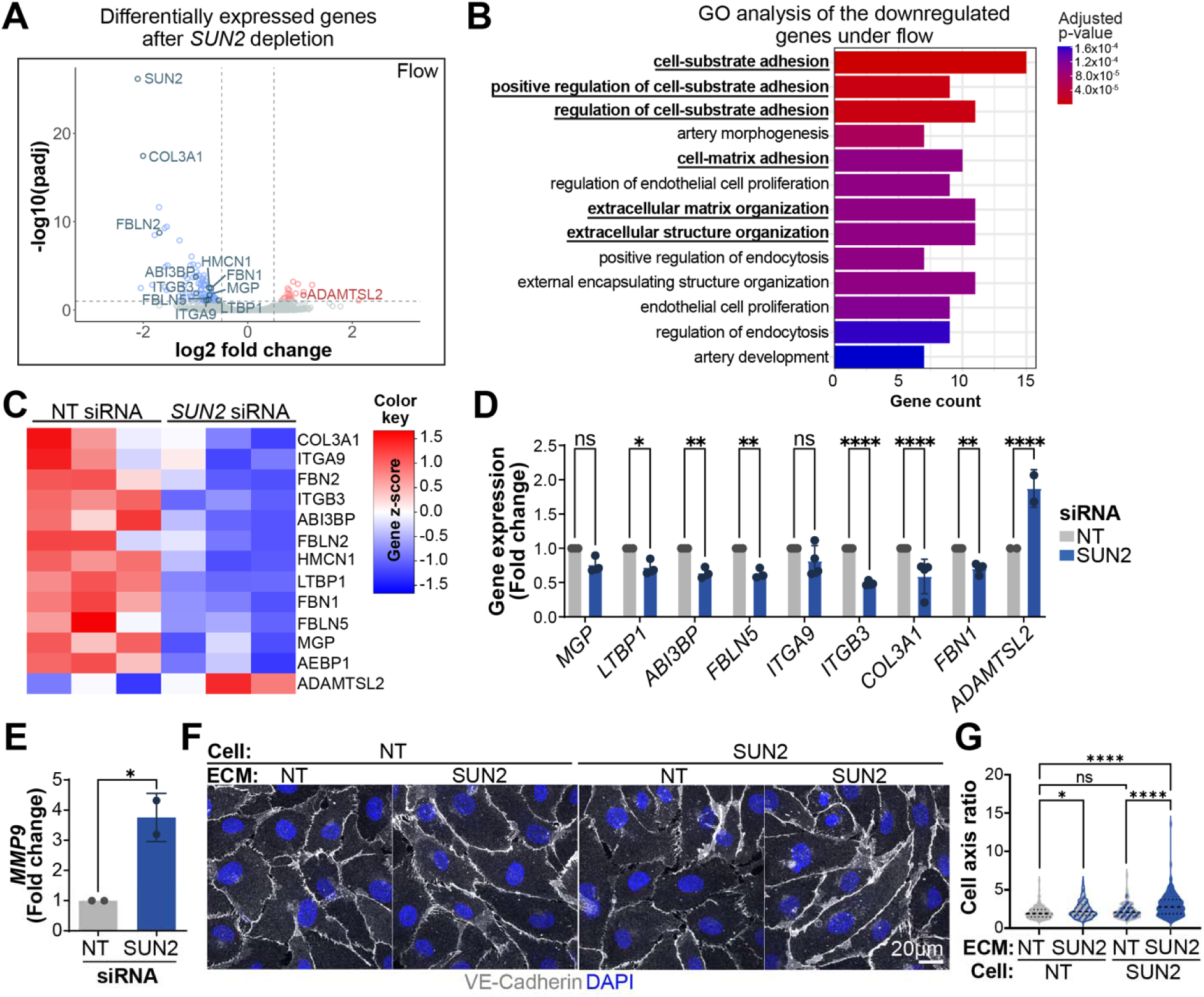
SUN2 regulates expression and function of endothelial ECM-adhesion-related components. **(A)** Volcano plot of differentially expressed genes from bulk RNA seq analysis of *SUN2*-depleted HUVEC exposed to laminar flow. Blue, downregulated genes; red, upregulated genes (n=3 biological replicates). **(B)** GO analysis of downregulated genes from bulk RNA seq data. **(C)** Relative expression levels of selected ECM-associated specific genes plotted using bulk RNA seq data (n=3 biological replicates). **(D, E)** RT-qPCR analysis of selected ECM-related genes (D) and *MMP9* (E) in HUVEC treated as indicated and exposed to laminar flow (n=2-3 biological replicates). **(F, G)** HUVEC treated with NT or SUN2 siRNA and seeded onto decellularized matrices derived from NT-or SUN2-depleted HUVEC for 4h. **(F)** Representative confocal images (maximum intensity projection) stained as indicated. Scale bar, 50 µm **(G)** Cell axis ratio quantification (n=3 biological replicates). Statistical analyses were performed using two-tailed Student’s t-tests **(D, E)** and one-way ANOVA with Tukey multiple comparisons tests **(G)**. ns, non-significant; *, p≤0.05; **, p ≤0.01; ***, p≤0.001; ****, p ≤ 0.0001.

This regulation was confirmed by RT-qPCR and included ECM genes encoding fibrillins (*FBN2* and, notably *FBN1*), collagen (*COL3A1*), fibulins (*FBLN2*, *FBLN5*), TGFβ-binding protein (*LTBP1*), and integrins (*ITGA9*, *ITGB3*) that participate in cell-matrix adhesion (**Fig. 2D)**. Interestingly, the subset of ECM-related genes regulated by SUN2 in endothelial cells under flow are important for large vessel integrity, and *FBN1* mutations lead to Marfan syndrome while *COL3A1* mutations are associated with vascular Ehlers-Danlos syndrome, both characterized by aneurysm and vessel rupture [55–57].

We next asked whether genes associated with ECM degradation were affected by *SUN2* depletion and did not see global mis-regulation of genes encoding these proteins (**Supp. Fig. 3B**) [58]. ADAMTSL2 expression is up-regulated under flow (**Fig. 2C-D)**, but this member of the ADAMTS (A Disintegrin And Metalloprotease with Thrombospondin Motifs) family lacks a metalloprotease domain [59] and thus cannot degrade matrix. Another class of ECM degradation genes, the matrix metalloproteases (MMPs), is associated with vessel wall weakness, dysfunction, and aneurysm [60].

Although MMPs were not globally mis-regulated by SUN2 depletion **(Supp. Fig. 3B)**, MMP9, a key regulator of ECM degradation [61], had increased RNA expression in SUN2-depleted cells compared to controls (**Fig. 2E**). Consistent with altered ECM composition, fibronectin protein levels were also reduced in SUN2-depleted EC (**Supp. Fig. 3C-D**). Taken together, the expression analysis shows that nuclear SUN2 depletion leads to down-regulation of genes encoding structural ECM components and increases expression of an ECM-degrading enzyme, suggesting that the matrix generated by SUN2-depleted endothelial cells is abnormal.

To test effects of *SUN2*-dependent matrix changes on endothelial cell behaviors, we performed ECM swap experiments, whereby control and *SUN2*-depleted cells were seeded on decellularized matrices derived from either condition for a short time **(Fig. 2F-G; Supp. Fig 3E**). *SUN2*-depleted cells seeded onto matrices from *SUN2*-depleted cells exhibited the expected abnormal hyper-elongation. Significantly, this phenotype was rescued to control levels in *SUN2*-depleted cells seeded onto matrices derived from control cells, indicating that ECM from SUN2-depleted cells differs from controls and that these changes contribute to endothelial cell phenotypes. Control cells seeded onto matrices from *SUN2*-depleted cells also exhibited limited but significant hyper-elongation. Thus, ECM from *SUN2*-depleted endothelial cells is altered, and these alterations, along with cell-based changes, contribute to the hyper-elongation phenotype of cells lacking SUN2.

### Nuclear SUN2 regulates localization and function of endothelial cell-matrix adhesions

Since SUN2-depleted endothelial cells cooperate with ECM to regulate cell elongation, we hypothesized that focal adhesions (FA), cellular structures on the basal surface that form contacts with matrix, are regulated by nuclear SUN2. Immunostaining for the FA components paxillin and vinculin revealed a significant increase in FA length in *SUN2*-depleted HUVEC, accompanied by a decrease in fluorescence intensity under flow or static conditions (**Fig. 3A-C; Supp. Fig. 4A-D**). Western blot analysis did not show significant differences in the expression of paxillin or vinculin (**Fig. 3D**), suggesting that SUN2 depletion alters FA dynamics rather than overall levels of FA components.

**Figure 3.**
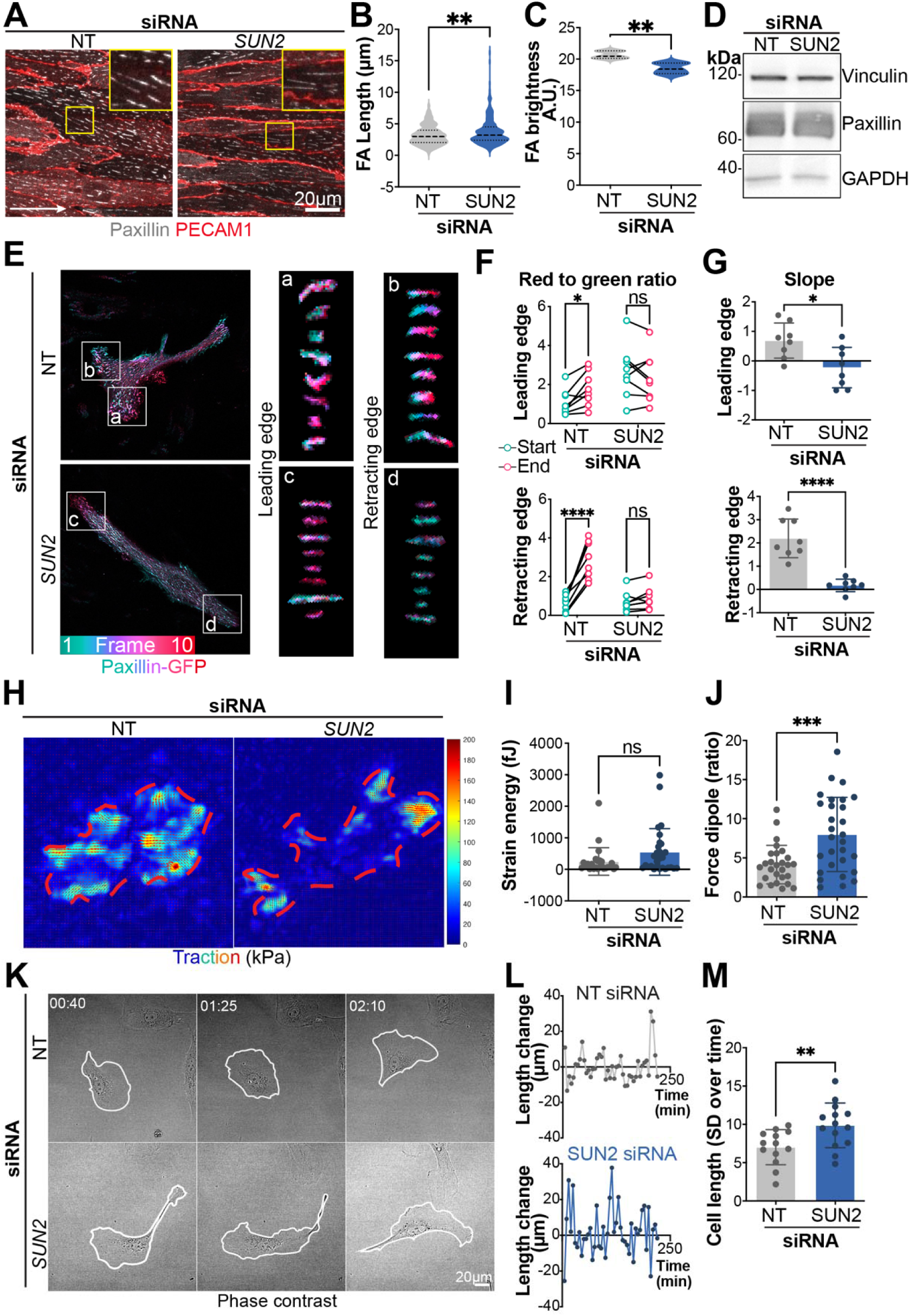
Nuclear SUN2 regulates the localization and function of endothelial cell-matrix adhesions. **(A)** Confocal images (maximum intensity projections) of HUVEC treated with siRNA and exposed to laminar flow, then fixed and stained as indicated. White arrow, flow direction. Scale bar, 20µm. **(B, C)** Quantification of indicated parameters (n=3 biological replicates). **(D)** Western blot analysis of vinculin, paxillin, and GAPDH expression in HUVEC treated with siRNA and exposed to laminar shear stress (15 dyn/cm^2^, 24h). **(E)** Color-coded temporal projections of paxillin–GFP expressing HUVEC treated with siRNA, with insets showing representative individual FA at the leading edge (a, c) and retracting edge (b, d). Images acquired at 4 min intervals over 40 min. **(F, G)** Quantification of FA dynamics in HUVEC treated with siRNA and expressing paxillin-GFP (n=3 biological replicates). **(F)** Red-to-green ratio quantification at the start and at the end of individual FA showing the temporal progression of the adhesion (time-color gradient) at the leading (top) and retracting (bottom) edges. **(G)** Slope quantification of the time-color gradient for individual FA at the leading (top) and retracting (bottom) edges. **(H)** Traction maps of HUVEC treated with siRNA on fibronectin-coated substrate. **(I, J)** Quantification of strain energy and force dipole ratio calculated from traction maps (n=3 biological replicates). **(K)** Representative phase-contrast images of migrating HUVEC treated with NT or SUN2 siRNA, at the indicated time points. **(L)** Quantification of major axis length over time, and **(M)** standard deviation of cell length showing increased variability in SUN2-depleted cells. Statistical analyses were performed using two-tailed Student’s t-tests, except for **(F),** which was analyzed using Multiple paired t-tests. ns, non-significant; *, p≤0.05; **, p ≤0.01; ***, p≤0.001.

Since both EC hyper-elongation **(Supp. Fig. 1D-E)** and FA changes induced by loss of SUN2 were not flow-dependent, subsequent functional analyses primarily utilized non-flow conditions. FA dynamics were analyzed using live imaging of paxillin-GFP or vinculin-mVenus (**Fig. 3E-G; Supp. Fig. 4E, Supp. Movies 1-2**). In control endothelial cells, FA exhibited continuous displacement over time, as indicated by a significant increase in the red-to-green ratio from the start to the end of the live-imaging period. In contrast, SUN2-depleted cells had significantly reduced FA displacement and reduced slope differential over the same time period, indicating reduced displacement and impaired FA dynamics. These results indicate that SUN2, located in the inner nuclear envelope, regulates focal adhesion organization and dynamics at the endothelial cell-matrix interface.

We next performed traction force microscopy on endothelial cells, an assay that measures the forces adherent cells use to deform an elastic substrate. We hypothesized that abnormal FA turnover would influence cell-matrix forces. *SUN2*-depleted cells showed similar overall strain energy compared to controls, but unlike controls with evenly distributed traction forces, *SUN2*-depleted cells had force-generating cell-matrix contacts concentrated at two distinct poles, leading to a significant change in the force dipole measurement (**Fig. 3H-J**). This finding is consistent with the lack of FA turnover at the extending edges of cells. We next asked whether the alterations in cell-substrate adhesion induced by *SUN2* loss affected cell migration. Consistent with altered traction forces, cell migration was impaired in SUN2-depleted cells, in line with previous studies associating increased force dipole ratio with non-migrating cells [62]. Scratch wound assays revealed a significant reduction in wound closure following *SUN2* loss, indicating defects in the coordinated migration of endothelial cells compared to controls (**Supp. Fig. 4H-I**). Endothelial cell migration at the single-cell level was also compromised. Controls exhibited directed migration, characterized by leading edge extension followed by coordinated retraction of the trailing edge; in contrast, SUN2-depleted cells often extended outward from both ends simultaneously, resulting in cell elongation and increased cell length, followed by abrupt cell retraction, as shown by increased variance in the length of the major cell axis over time (**Fig. 3K-M; Supp. Movies 3-4**). Thus, nuclear SUN2 is essential for proper endothelial cell migration.

### Nuclear SUN2 regulates microtubule stability in endothelial cells and utilizes nesprin-1

SUN2 regulates ECM properties and cell-matrix interactions, cellular processes that occur far from the nucleus. Our previous work revealed that LINC complex interactions involving SUN1 regulate microtubule dynamics to affect endothelial cell-cell adhesions [32], and microtubules influence cell polarity and migration by regulating focal adhesion turnover [63–65]. Thus, we hypothesized that SUN2 regulates the microtubule cytoskeleton, and we found that both microtubule density and acetylated tubulin levels, a marker of microtubule stabilization, were significantly increased in *SUN2*-depleted endothelial cells compared to controls (**Fig. 4A-D**). To assess microtubule dynamics, we performed live image tip-tracking of labeled microtubule plus ends and found that microtubules spent more time growing and less time shrinking in SUN2-depleted endothelial cells compared to controls **(Fig. 4E-F; Supp. Movies 5-6)**. Taken together, these findings indicate that microtubule over-stabilization downstream of nuclear SUN2 perturbation correlates with endothelial cell dysfunction.

**Figure 4.**
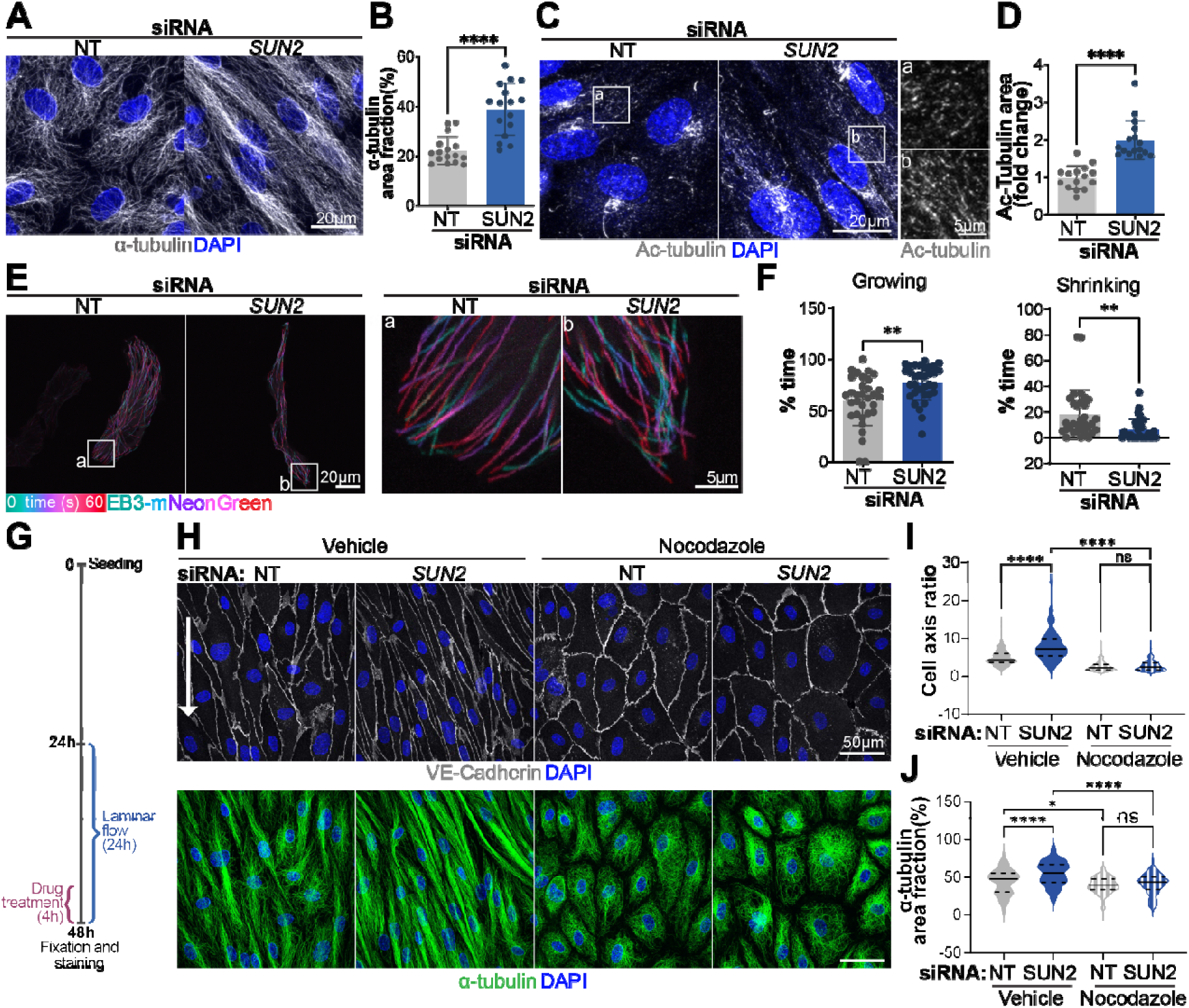
Nuclear SUN2 destabilizes microtubules in endothelial cells. (A, C) Confocal images (maximum intensity projections) of HUVEC treated with siRNA, fixed and stained as indicated. Scale bar, 20µm. **(C: a-b)** Acetylated α-tubulin staining shown at higher magnification. Scale bar, 5µm. **(B, D)** Quantification of indicated parameters. (n = 3 replicates). **(E)** Temporal projection of EB3-mNeonGreen expressing HUVEC treated with siRNA as indicated. Images acquired at 470ms intervals over 2 min. **(F)** Quantification of microtubule dynamics using the indicated parameters. (n= 10 microtubules from 3 different cells). **(G)** Diagram of nocodazole treatment under orbital flow conditions. **(H)** Confocal images (maximum intensity projections) of HUVEC with the indicated siRNAs, exposed to laminar orbital flow for 24h and treated with DMSO (vehicle) or nocodazole for the last 4h, followed by fixation and staining as indicated. Scale bar: 20µm. VE-Cad; VE-Cadherin. **(I, J)** Quantification of indicated parameters (n=3 replicates). Statistical analyses were performed using two-tailed Student’s t-test, except for **(I, J),** which was analyzed using one-way ANOVA with Tukey multiple comparisons tests. *, p ≤0.05; **, p≤ 0.01 ****, p≤ 0.0001; ns, not significant.

To determine whether microtubules are required for endothelial cell perturbations induced by *SUN2* loss, we partially depolymerized the microtubule cytoskeleton via low-dose nocodazole exposure and examined the effect on cell shape. As predicted, this treatment rescued the increased microtubule density and the hyper-elongation phenotype of SUN2-depleted cells under laminar shear stress (**Fig. 4G-J**). Thus, SUN2 regulates endothelial cell responses to environmental cues by controlling cell shape and flow responses, likely through effects on microtubule organization and dynamics.

The LINC complex interaction with the microtubule cytoskeleton is mediated via nesprins. We previously showed that SUN1 regulates the microtubule cytoskeleton in endothelial cells via its interaction with nesprin-1 (encoded by *SYNE1*) [32], so we hypothesized that SUN2-mediated effects on the microtubule network also require nesprin-1. Although nesprin-1 depletion alone did not affect either the cell axis ratio or microtubule density, co-depletion of SUN2 and nesprin-1 rescued both the hyper-elongation and the increased microtubule density induced by SUN2 loss alone (**Fig. 5A-C)**. These results suggest that SUN2 controls hyper-elongation and microtubule stability in endothelial cells through a nesprin-1-dependent mechanism.

**Figure 5.**
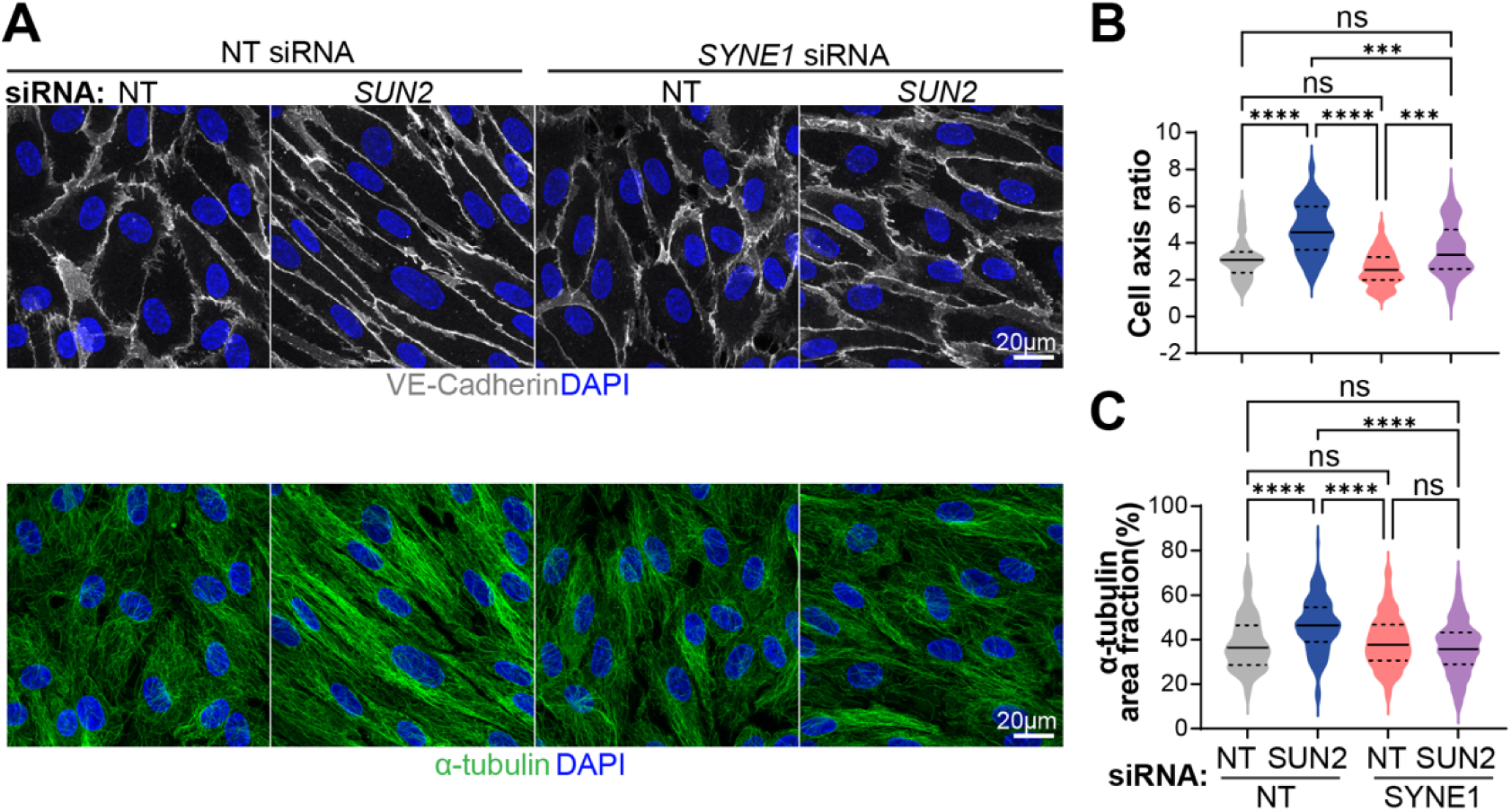
Nesprin-1 is required for SUN2-mediated effects on the microtubule cytoskeleton. **(A)** Confocal images (maximum intensity projections) of HUVEC with the indicated siRNAs, treated with siRNA, fixed, and stained as indicated. Scale bar, 20µm. **(B, C)** Quantification of indicated parameters (n=3 biological replicates). Statistical analyses were performed using one-way ANOVA with Tukey multiple comparisons tests. ***, p ≤ 0.001; ****, p ≤ 0.0001; ns, not significant.

### Nuclear SUN2 is required for proper endothelial barrier function

Endothelial cell-matrix adhesion is essential for barrier function [66]. Since *SUN2* depletion perturbs cell-matrix interactions and leads to significant downregulation of *ITGB3* expression (**Fig. 2C-D**), a gene required for vascular barrier function [67], we asked whether nuclear SUN2 regulates endothelial barrier integrity. Using real-time cell analysis (RTCA) to measure electrical impedance across endothelial cell monolayers, we found that *SUN2*-depleted cells had a reduced cell index throughout a 24h time course (**Fig. 6A-B)**, indicating impaired monolayer integrity. Consistent with this result, a functional assay based on streptavidin accessibility to biotinylated matrix [32, 43, 44] revealed increased labeling following *SUN2* depletion (**Fig. 6C-D, Supp. Fig. 5A**).

**Figure 6.**
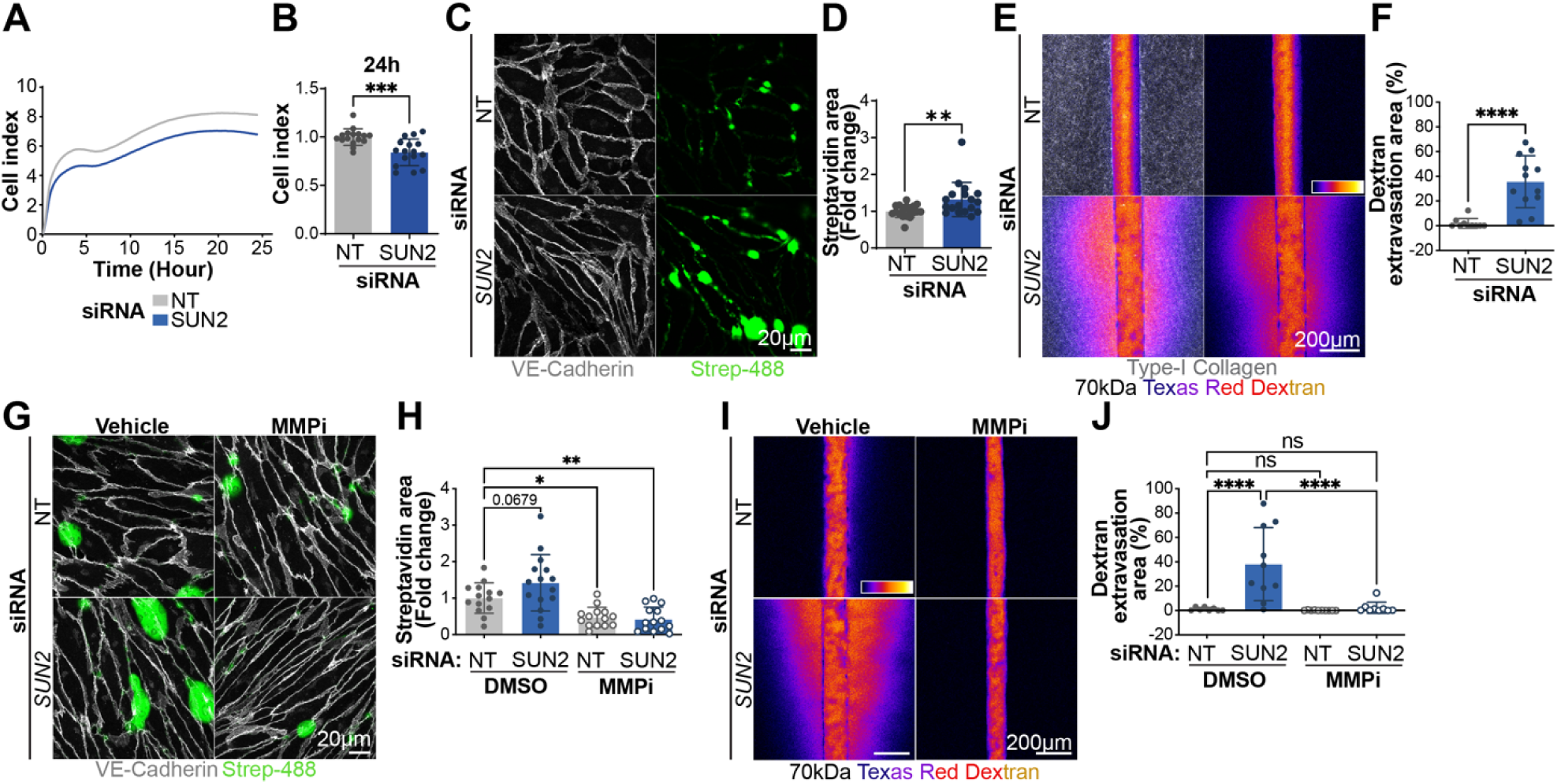
Nuclear SUN2 is required for proper endothelial barrier function. **(A)** Representative curve of cell index measured by real-time cell analysis (RTCA) in HUVEC treated with siRNA as indicated. **(B)** Quantification of cell index measured by RTCA at 24h. (n=4 biological replicates). **(C)** Confocal images (maximum intensity projections) of HUVEC with indicated siRNAs, on biotinylated-fibronectin, treated with streptavidin-488 (Strep-488), and stained as indicated. Scale bar, 20µm. **(D)** Quantification of streptavidin area in HUVEC with indicated siRNAs. (n=5 biological replicates). **(E)** Representative images of 3D vessels seeded with HUVEC treated with siRNA as indicated. 70kDa Texas Red-dextran (Fire-LUT) was injected into the channel, and leakage was monitored over time. Left panels: type-I collagen matrix and dextran fluorescence at 02:30 min. Right panels: dextran fluorescence at 02:30min. **(F)** Quantification of dextran extravasation area (% of total extravascular area) at the 02:30 min time point (n=3 biological replicates). **(G)** Representative confocal images (maximum intensity projections) of HUVEC with indicated siRNAs, plated on biotinylated-fibronectin and treated as indicated for 24h, followed by incubation with streptavidin-488 (Strep-488), fixation, and staining as indicated. Scale bar: 20µm. **(H)** Quantification of (G). (n=3 biological replicates). **(I)** Representative images of 3D blood vessels with indicated siRNAs and treatments. Dextran fluorescence at 02:30min. **(J)** Quantification of (I) (n=3 biological replicates). Statistical analyses were performed using two-tailed Student’s t-tests (B, D, F) and one-way ANOVA with Tukey multiple comparison tests (H, J). *, p≤0.05; **, p ≤0.01; ***, p≤0.001; ****, p ≤ 0.0001; ns, not significant.

Endothelial barrier function was further evaluated using 3D micro-fabricated vessels exposed to a 70-kDa fluorescent dextran in the lumen. Live imaging revealed that vessels formed with *SUN2*-depleted endothelial cells had significant dextran leakage compared to controls (**Fig. 6E-F**). Taken together, these findings show that SUN2 regulates endothelial cell monolayer integrity and barrier function. Because SUN2 depletion leads to elevated *MMP9* mRNA expression, induces ECM remodeling, and over-stabilizes microtubules that deliver MMPs, we asked whether excessive MMP activity contributes to the observed barrier defects. Inhibition of MMP activity using the broad-spectrum inhibitors Marimastat or Ilomastat rescued the increased streptavidin labeling induced by SUN2 depletion in 2D (**Fig. 6G-H, Supp. Fig. 5B-C**). In addition, Marimastat treatment rescued dextran extravasation in the 3D micro-fabricated vessels seeded with SUN2-depleted cells (**Fig. 6I-J**). Thus, SUN2 is essential for maintaining endothelial barrier integrity, likely by regulating MMP expression and microtubule-dependent trafficking to the ECM.

## DISCUSSION

Molecular complexes that reside in the nuclear envelope interact with cellular structures to regulate cell behaviors far from the nucleus. Here, we show that SUN2, a component of the nuclear LINC complex, regulates the responses of vascular endothelial cells to environmental cues and cell-matrix interactions in cultured cells and *in vivo*. Moreover, we find that SUN2 influences the properties of both the secreted matrix and focal adhesions that mediate these interactions, likely via dynamic regulation of the microtubule cytoskeleton **(Fig. 7)**. These interconnected mechanisms regulate vascular permeability, and SUN2 perturbation leads to alterations in vascular matrix organization and barrier integrity that resemble those observed in vessel wall diseases such as Marfan syndrome. Thus, nuclear SUN2 regulates cell-matrix interactions that extend to the surrounding microenvironment and are essential for vascular function.

**Figure 7.**
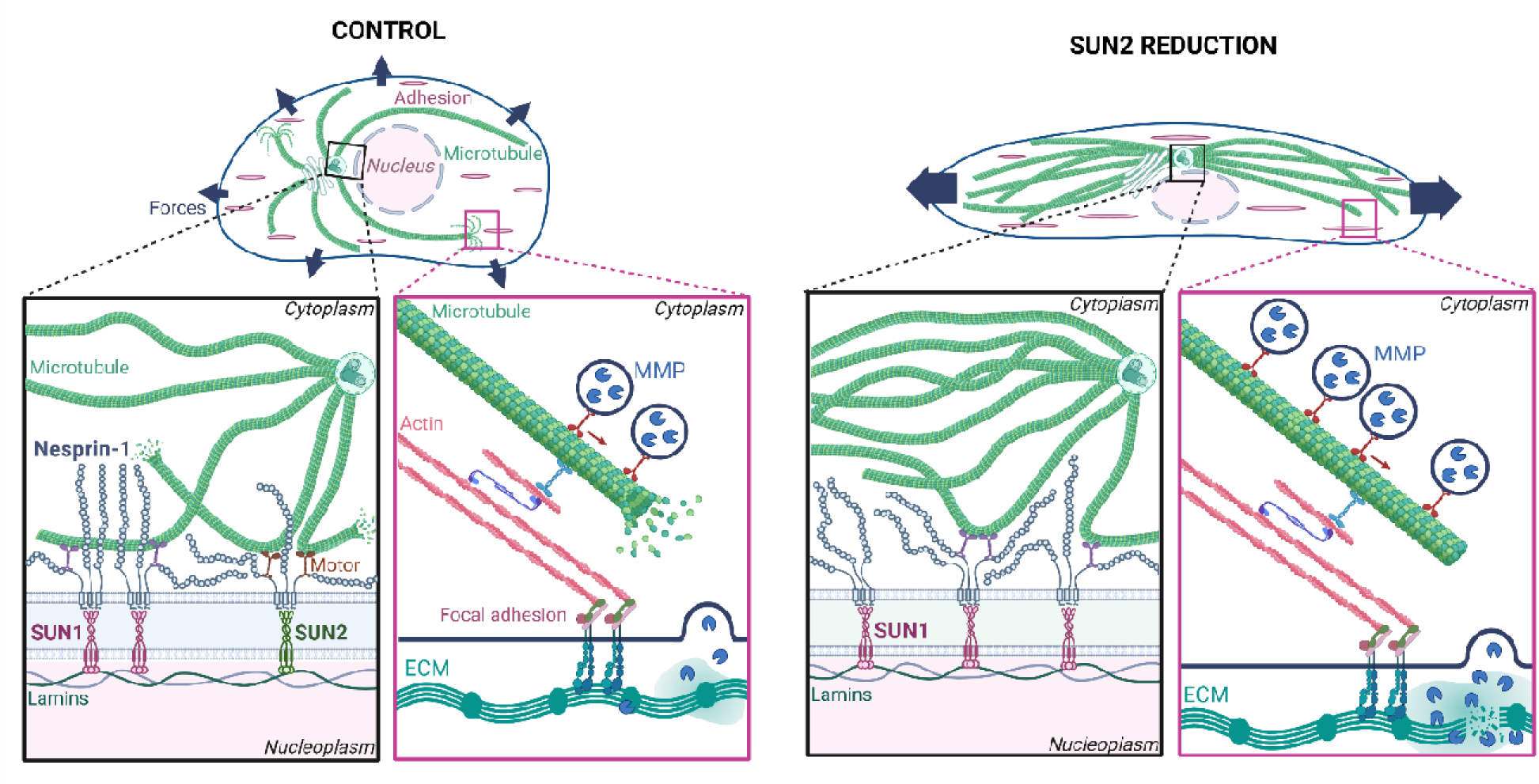
Proposed model for SUN2 regulation of endothelial cell-matrix interactions. In control endothelial cells (left panel), a balanced distribution of SUN1/nesprin-1 and SUN2/nesprin-1 LINC complexes at the nuclear envelope supports proper microtubule dynamics (black square), leading to regulated delivery of MMPs to the ECM (pink square). In contrast, SUN2-depletion disrupts LINC complex balance, leading to increased microtubule stability (black square) and enhanced delivery of MMPs to the ECM (pink square).

SUN1 and SUN2 form distinct homo-trimeric LINC complexes with nesprins that integrate with and regulate cellular processes [68]. We previously showed that SUN1 regulates endothelial cell-cell junction integrity, although its loss does not alter endothelial transcriptional profiles [32]. In contrast, SUN2 depletion led to down-regulation of numerous genes in endothelial cells, including a subset of genes encoding ECM proteins that contribute to the integrity and elasticity of large vessel walls, such as *COL3A1*, *FBN1*, and *ITGB3* [67, 69, 70]. The gene expression changes caused by endothelial SUN2 loss were accompanied by modifications in elastin profiles and junction morphology *in vivo*, changes that are characteristic of Marfan syndrome, a connective tissue disorder linked to reduced function FBN1 mutations that result in large vessel aneurysms [49, 71]. Our endothelial cell-ECM swap experiments revealed that SUN2-depleted endothelial cells initially had a normal cell axis ratio on control ECM. This is consistent with findings that vessel wall matrix properties influence multiple cell parameters, including adhesion, migration, and mechanotransduction, and contribute to vascular diseases [72–74]. It is also consistent with the finding that global *Sun2* loss is associated with a blunted fibrotic response in cardiomyocytes (Stewart et al, 2019).

The swap experiments showed that matrix defects alone did not lead to endothelial cell changes downstream of SUN2 loss, suggesting additional cellular adaptations. These alterations prominently affected focal adhesion dynamics, with SUN2-depleted cells showing reduced focal adhesion turnover and spatially mis-placed force-generating contacts between EC and the matrix, resulting in altered cell migration. Interestingly, similar defects were observed in HUVEC following nesprin-1 loss [27], indicating that the nucleus functions as a central hub for force coordination in cells through the LINC complex. Microtubules regulate focal adhesion turnover via dynamic interactions [65, 75], and we found that SUN2-depleted EC have over-stabilized microtubules with altered growth and retraction dynamics. Although pharmacological stabilization of microtubules prevents flow-mediated alignment in EC [15, 76], SUN2-depleted endothelial cells have an abnormally increased cell-axis ratio under both static and flow conditions, suggesting that the microtubule over-stabilization induced by SUN2 loss is more subtle. Taken together, our findings indicate that loss of SUN2 impairs the integration and transmission of incoming mechanical signals. This defect in turn is predicted to result in impaired flow responses and migration, consistent with an inability to properly set cell polarity.

Vascular permeability is impaired in SUN2-depleted EC. This dysfunction correlates with increased expression of matrix-degrading MMP9 and is rescued by blockade of MMP activity, suggesting that barrier dysfunction results from aberrant matrix degradation. MMP delivery to the matrix is thought to be increased by microtubule residence at focal adhesions [77, 78], consistent with our finding that MTs are over-stabilized in SUN2-deficient cells. Thus, although changes in ECM-related gene expression likely contribute to matrix defects, increased MMP-mediated degradation is crucial for matrix changes leading to barrier dysfunction. Moreover, changes in matrix properties and degradation are hallmarks of diseases that compromise vessel wall integrity, including Marfan syndrome [79, 80].

Our data indicate that SUN2 regulates the microtubule cytoskeleton in a manner opposite to that of SUN1, with SUN1 stabilizing and SUN2 destabilizing microtubule dynamics. These findings are consistent with a model linking LINC complex function at the nuclear membrane to events at cell-cell and cell-matrix junctions in endothelial cells (**Fig. 7)**. Reports suggesting that SUN1 and SUN2 LINC complexes differ in their outputs and that SUN1 can antagonize or balance SUN2 complexes support this model [81, 82]. We previously hypothesized that SUN1 sequesters nesprin-1 from a second nuclear membrane component [32], and our findings here suggest that SUN2 is that component. However, SUN1/nesprin-1 complexes also likely function, as the detrimental effects of SUN2 loss require nesprin-1. In this model, SUN2/nesprin-1 complexes confer an important polarization/force distribution function to endothelial cells and SUN1/nesprin-1 complexes that, when lost, lead to dysfunctional cell-matrix interactions. Conversely, when SUN2/nesprin-1 complexes are more abundant than SUN1/nesprin-1 complexes, they cause destabilization of peripheral microtubules, leading to increased contractility driven by elevated localized Rho signaling [32]. For these LINC complex functions, nesprin-1 may be rate-limiting and drive the balance of SUN1/nesprin-1 vs. SUN2/nesprin-1 complexes in the nuclear envelope that normally regulate microtubules. This model is consistent with our finding that nesprin-1 is required for the adverse effects of loss of either SUN protein on EC functions. Our data show that over-stabilized microtubules more profoundly affect endothelial cell-matrix interactions and flow alignment, while destabilized microtubules more profoundly affect contractility and cell-cell integrity, and both disturbances lead to loss of barrier function.

One important limitation of our study is that the sequence of events leading to downstream changes in microtubule dynamics following selective LINC complex disruption is not known. Since LINC complexes also interact with and regulate the actin cytoskeleton, it is possible that LINC complexes directly affect actin dynamics that then feedback on microtubules. Alternatively, SUN1/nesprin-1 and SUN2/nesprin-1 LINC complexes may differentially interact with microtubule motors such as kinesins. These plus-end-directed motors bind nesprins via a motif close to the transmembrane domain on the cytoplasmic side [83, 84], and they can process towards the microtubule plus end with the nucleus as cargo. Motors also affect microtubule stability in context-dependent ways, and both stabilization and de-stabilization effects of motor procession have been reported [85]. Thus, selective binding or movement of motors may differentially affect microtubules downstream of SUN1 vs. SUN2 LINC complexes. In summary, our work shows that SUN2 has distinct roles from SUN1 in vascular endothelial cells and blood vessels, that these roles converge on endothelial cell responses to their environment and differentially regulate microtubule dynamics, and that endothelial SUN2 disruption changes cell-matrix interactions in ways consistent with vessel wall disease, suggesting new therapeutic targets for these diseases.

## SUPPLEMENTAL FIGURE LEGENDS

**Supplementary Figure 1 (for Fig 1). SUN2 regulates endothelial cell axis ratio and measurements. (A)** Confocal images (maximum intensity projections) of HUVEC with indicated siRNA and stained as indicated. Scale bar, 20µm. **(B)** Quantification of SUN2 nuclear intensity in HUVEC with indicated siRNA. **(C)** Diagram illustrating cell axis ratio quantification. **(D)** Confocal images (maximum intensity projections) of HUVEC with indicated siRNA and stained as indicated, and **(E)** cell axis ratio quantification (n=3 biological replicates). **(F)** Schematic illustrating cell polarity quantification, based on nucleus and Golgi staining. **(G)** Schematic showing the breeding strategy to generate *Sun2^iECKO^* mice. **(H)** Schematic illustrating EC junction measurement quantification. Statistical analyses: two-tailed Student’s t-tests. ****, p≤0.0001.

**Supplementary Figure 2 (for Fig 1). Nuclear SUN2 regulates vascular development. (A)** Representative confocal images (maximum intensity projections) of P7 mouse retinas from indicated genotypes, stained as indicated. Scale bar, 200 µm. **(B, C)** Quantification of the indicated parameters (WT, n=5, Cdh5-Cre/+, n=5, *Sun2^fl/fl^*, n=11; *Sun2^iECKO^,* n=9). **(D, E)** Confocal images (*Tg(fli:LifeAct-GFP)* and quantification of zebrafish embryos with indicated morpholino (MO) treatments at 36hpf. Scale bar: 100 µm. (NT MO, n=22; *sun2* MO, n=22). **(F, G)** Confocal images (*Tg(fli:LifeAct-GFP)* and quantification of zebrafish embryos with indicated genotypes at 36hpf. Scale bar, 100 µm. (wt, n=5; *sun2^-/-^*, n=4). Statistical analyses were performed using two-tailed Student’s t-tests, except for **(B),** which was analyzed using one-way ANOVA with Tukey multiple comparisons test. *, p≤0.05; ***, p≤0.001; ****, p≤0.0001; ns, not significant.

**Supplementary Figure 3 (for Fig 2). SUN2 regulates extracellular matrix gene expression and composition. (A, B)** Relative expression levels of ECM-associated genes and ECM-degradation associated genes plotted using bulk RNA seq data of HUVEC treated as indicated and cultured under laminar shear stress (n=3 biological replicates). **(C, D)** Western blot analysis of fibronectin and GAPDH expression in HUVEC treated with siRNA, seeded onto gelatin matrix, and exposed to laminar shear stress (15 dyn/cm^2^, 24h) (n=3 biological replicates). **(E)** Diagram of matrix swap experiments.

**Supplementary Figure 4 (for Fig.3)**. **Nuclear SUN2 regulates focal adhesion dynamics and collective cell migration in endothelial cells. (A)** Confocal images (maximum intensity projections) of HUVEC treated with siRNA, cultured under static conditions, then fixed and stained for paxillin. Scale bar, 20µm. **(B)** Quantification of indicated parameters (n=3 biological replicates). **(C)** Confocal images (maximum intensity projections) of HUVEC treated with siRNA, cultured under static conditions, then fixed and stained for vinculin. Scale bar, 20µm. **(D)** Quantification of indicated parameters (n=3 biological replicates). **(E)** Temporal projections of vinculin–mVenus expressing HUVEC treated with siRNA, with insets showing representative individual FA at the leading edge (a,c) and retracting edge (b,d). Images were acquired at 4 min intervals over 40 min. **(F, G)** Quantification of FA dynamics in vinculin-mVenus expressing HUVEC treated with siRNA (n=2 biological replicates). **(F)** Red-to-green ratio quantification at the start and at the end of individual FA showing the temporal progression of the adhesion (time-color gradient) at the leading (top) and retracting (bottom) edges. **(G)** Slope quantification of the time-color gradient for individual FA at the leading (top) and retracting (bottom) edges. **(H)** Scratch wound healing assay of HUVEC treated with siRNA as indicated, and **(I)** quantification of the recovered area after 8h (n=5 replicates). Statistical analyses were performed using two-tailed Student’s t-tests, except for (F), which was analyzed using Multiple paired t-tests. ns, non-significant; *, p≤0.05; **, p ≤0.01; ***, p≤0.001; ****, p≤0.0001.

**Supplementary Figure 5 (for Fig.6). Nuclear SUN2 regulates endothelial barrier function via MMP activity. (A)** Schematic illustrating the biotin-streptavidin labeling assay to assess monolayer integrity. **(B)** Confocal images (maximum intensity projections) of HUVEC with indicated siRNAs, on biotinylated-fibronectin, and treated as indicated for 24h, followed by incubation with Strep-488, fixation, and staining as indicated. Scale bar, 20µm. **(C)** Quantification of streptavidin area in HUVEC with indicated siRNAs and treatments (n=3 biological replicates).

## SUPPLEMENTAL MOVIE LEGENDS

**Supplementary Movie 1 (for Fig. 3E). Focal adhesion dynamics in control endothelial cells.** Movies acquired by confocal imaging of paxillin–GFP expressing HUVEC treated with NT siRNA. Images acquired at 4 min intervals over 40 min. Scale bar: 5µm.

**Supplementary Movie 2 (for Fig. 3E). SUN2 depletion affects focal adhesion dynamics in endothelial cells.** Movies acquired by confocal imaging of paxillin–GFP expressing HUVEC treated with SUN2 siRNA. Images acquired at 4 min intervals over 40 min. Scale bar: 5µm.

**Supplementary Movie 3 (for Fig. 3K). Coordinated migration of control endothelial cells.** Movies acquired by phase-contrast confocal imaging of HUVEC treated with NT siRNA. Images acquired at 5 min intervals over 235 min. Scale bar: 20µm.

**Supplementary Movie 4 (for Fig. 3K). SUN2 depletion disrupts coordinated migration of endothelial cells.** Movies acquired by phase-contrast confocal imaging of HUVEC treated with SUN2 siRNA. Images acquired at 5 min intervals over 235 min. Scale bar: 20µm.

**Supplementary Movie 5 (for Fig. 4E). Microtubule dynamics in control endothelial cells.** Movies acquired using spinning disc microscopy on EB3-mNeonGreen expressing HUVEC treated with NT siRNA. Images acquired at 470ms intervals over 60 sec. Black arrows indicate shrinking microtubules. Scale bar: 5µm.

**Supplementary Movie 6 (for Fig. 4E). SUN2 depletion increases microtubule stability in endothelial cells.** Movies acquired using spinning disc microscopy on EB3-mNeonGreen expressing HUVEC treated with SUN2 siRNA. Images acquired at 470ms intervals over 60 sec. Black arrows indicate shrinking microtubules. Scale bar: 5µm.

## Notes

FUNDING This work was supported by a grant from NIH-NHLBI (R35HL139950 to VLB), an American Heart Association Postdoctoral Fellowship (25POST1377401 to PB), and grants from NIH-NIGMS (R35 GM142944 to WJP and MR, R35GM158040 to WRL), American Heart Association Predoctoral (19PRE34380887 to DBB) and Postdoctoral (AHA829371 to MO) Fellowships, and a Ruth L. Kirschstein Predoctoral Fellowship (1F31HL156527-01 to NT).

### Competing Interest Statement

The authors have declared no competing interest.

