## Supplementary Material for "Nuclear SUN2 coordinates endothelial cell-matrix interactions to regulate blood vessel homeostasis and barrier function"

**Supplemental Figures:** Pages 2-7

**Supplemental Movie Legends:** Page 8

**Supplemental Tables:** Pages 9-12

### SUPPLEMENTAL FIGURES

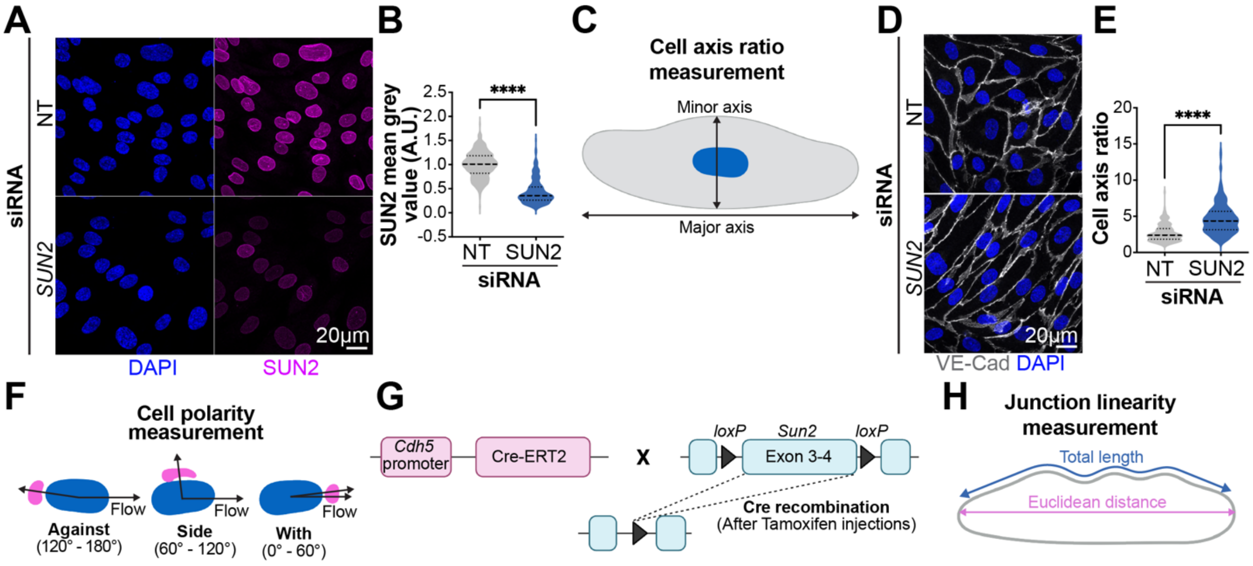


**Supplementary Figure 1 (For Fig 1). SUN2 regulates endothelial cell axis ratio and measurements. (A)** Confocal images (maximum intensity projections) of HUVEC with indicated siRNA and stained as indicated. Scale bar, 20µm. **(B)** Quantification of SUN2 nuclear intensity in HUVEC with indicated siRNA. **(C)** Diagram illustrating cell axis ratio quantification. **(D)** Confocal images (maximum intensity projections) of HUVEC with indicated siRNA and stained as indicated, and **(E)** cell axis ratio quantification (n=3 biological replicates). **(F)** Schematic illustrating cell polarity quantification, based on nucleus and Golgi staining. **(G)** Schematic showing the breeding strategy to generate *Sun2*$^{iECKO}$ mice. **(H)** Schematic illustrating EC junction measurement quantification. Statistical analyses: two-tailed Student's t-tests**.** ****, p≤0.0001.

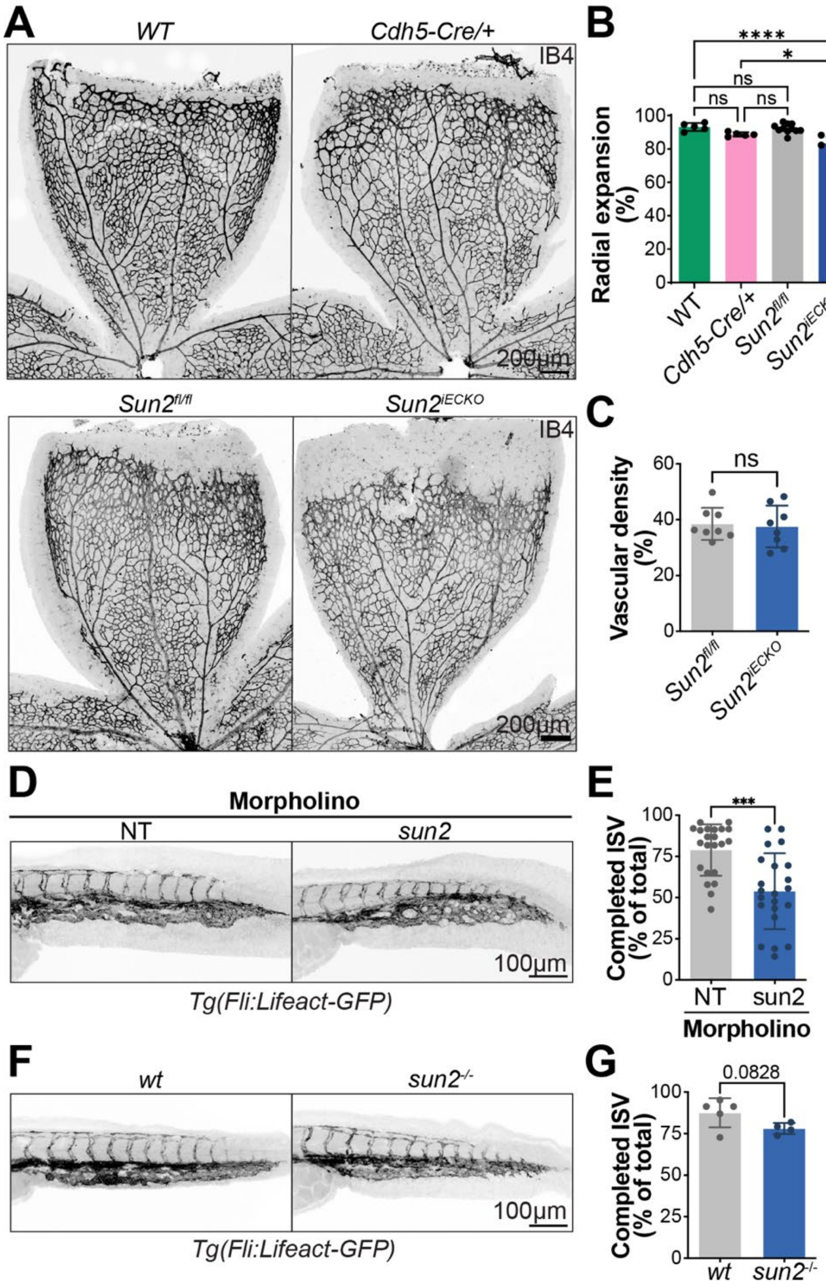


**Supplementary Figure 2 (For Fig 1). Nuclear SUN2 regulates vascular development. (A)** Representative confocal images (maximum intensity projections) of P7 mouse retinas from indicated genotypes, stained as indicated. Scale bar, 200 µm. **(B, C)** Quantification of the indicated parameters (WT, n=5, Cdh5-Cre/+, n=5, *Sun2*$^{fl/fl}$, n=11; *Sun2*$^{iECKO}$, n=9). **(D, E)** Confocal images (*Tg(fli:LifeAct-GFP)* and quantification of zebrafish embryos with indicated morpholino (MO) treatments at 36hpf. Scale bar: 100 µm. (NT MO, n=22; *sun2* MO, n=22). **(F, G)** Confocal images (*Tg(fli:LifeAct-GFP)* and quantification of zebrafish embryos with indicated genotypes at 36hpf. Scale bar, 100 µm. (wt, n=5; *sun2*$^{-/-}$, n=4). Statistical analyses were performed using two-tailed Student's t-tests, except for **(B),** which was analyzed using one- way ANOVA with Tukey multiple comparisons test. *, p≤0.05; ***, p≤0.001; ****, p≤0.0001; ns, not significant.

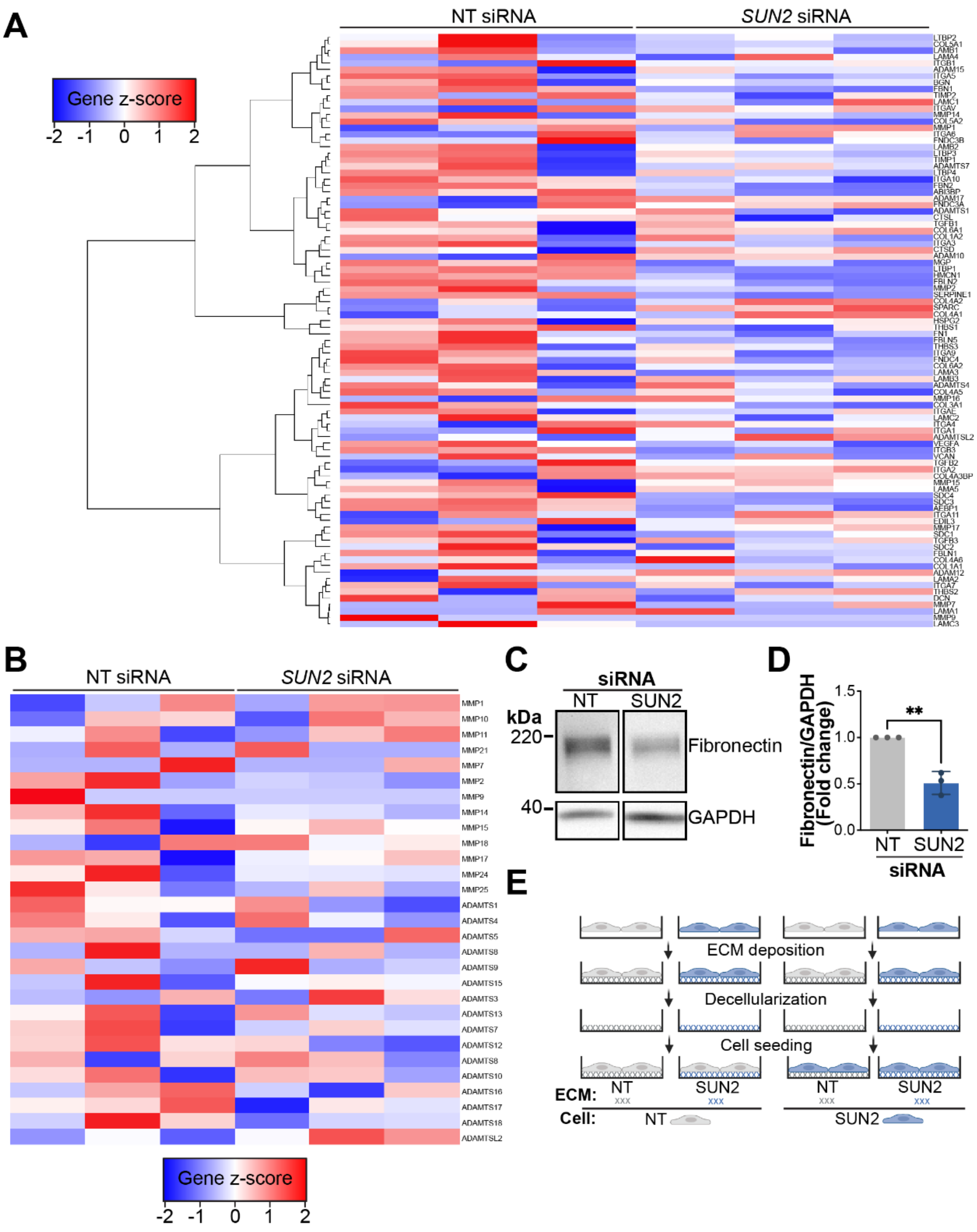


**Supplementary Figure 3 (For Fig 2). SUN2 regulates extracellular matrix gene expression and composition.** **(A, B)** Relative expression levels of ECM-associated genes and ECM-degradation associated genes plotted using bulk RNA seq data of HUVEC treated as indicated and cultured under laminar shear stress (n=3 replicates). **(C, D)** Western blot analysis of fibronectin and GAPDH expression in HUVEC treated with siRNA, seeded onto gelatin matrix, and exposed to laminar shear stress (15 dyn/cm$^2$, 24h) (n=3 biological replicates). **(E)** Diagram of matrix swap experiments.

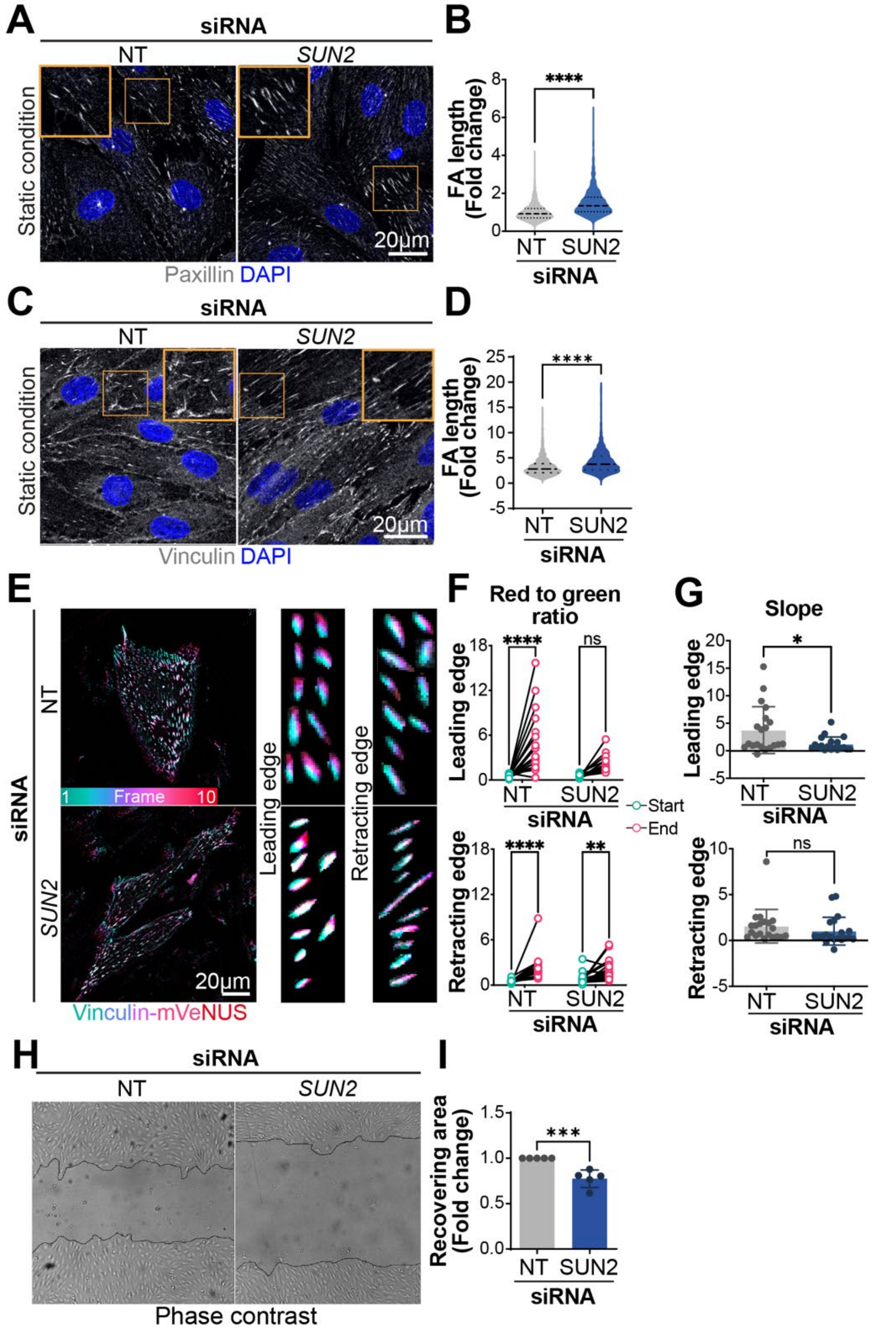


**Supplementary Figure 4 (For Fig.3)**. **Nuclear SUN2 regulates focal adhesion dynamics and collective cell migration in endothelial cells. (A)** Confocal images (maximum intensity projections) of HUVEC treated with siRNA, cultured under static conditions, then fixed and stained for paxillin. Scale bar, 20µm. **(B)** Quantification of indicated parameters (n=3 biological replicates). **(C)** Confocal images (maximum intensity projections) of HUVEC treated with siRNA, cultured under static conditions, then fixed and stained for vinculin. Scale bar, 20µm. **(D)** Quantification of indicated parameters (n=3 biological replicates). **(E)** Temporal projections of vinculin–mVenus expressing HUVEC treated with siRNA, with insets showing representative individual FA at the leading edge (a,c) and retracting edge (b,d). Images were acquired at 4 min intervals over 40 min. **(F, G)** Quantification of FA dynamics in vinculin-mVenus expressing HUVEC treated with siRNA (n=2 biological replicates). **(F)** Green-to-red ratio quantification at the start and at the end of individual FA showing the temporal progression of the adhesion (time-color gradient) at the leading (top) and retracting (bottom) edges. **(G)** Slope quantification of the time-color gradient for individual FA at the leading (top) and retracting (bottom) edges. **(H)** Scratch wound healing assay of HUVEC treated with siRNA as indicated, and **(I)** quantification of the recovered area after 8h (n=5 biological replicates). Statistical analyses were performed using two-tailed Student's t-tests, except for **(F),** which was analyzed using Multiple paired t-tests**.** ns, non-significant; *, $p \leq 0.05$; **, $p \leq 0.01$; ***, $p \leq 0.001$; ****, $p \leq 0.0001$.

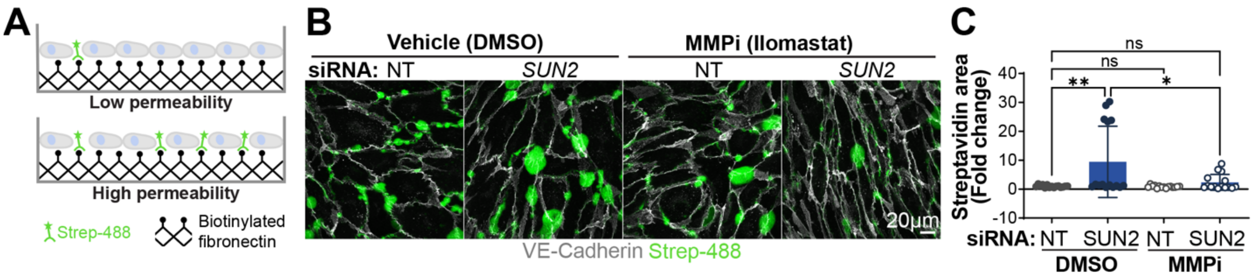


**Supplementary Figure 5 (For Fig.6). Nuclear SUN2 regulates endothelial barrier function via MMP activity. (A)** Schematic illustrating biotin-streptavidin labeling assay to assess monolayer integrity. **(B)** Confocal images (maximum intensity projections) of HUVEC with indicated siRNAs, on biotinylated-fibronectin, and treated as indicated for 24h, followed by incubation with Strep-488, fixation, and staining as indicated. Scale bar, 20µm. **(C)** Quantification of streptavidin area in HUVEC with indicated siRNAs and treatments (n=3 biological replicates).

### SUPPLEMENTAL MOVIE LEGENDS

**Supplementary Movie 1 (For Fig. 3E). Focal adhesion dynamics in control endothelial cells.** Movies acquired by confocal imaging of paxillin–GFP expressing HUVEC treated with NT siRNA. Images acquired at 4 min intervals over 40 min. Scale bar: 5μm.

**Supplementary Movie 2 (For Fig. 3E). SUN2 depletion affects focal adhesion dynamics in endothelial cells.** Movies acquired by confocal imaging of paxillin–GFP expressing HUVEC treated with SUN2 siRNA. Images acquired at 4 min intervals over 40 min. Scale bar: 5μm.

**Supplementary Movie 3 (For Fig. 3K). Coordinated migration of control endothelial cells.** Movies acquired by phase-contrast confocal imaging of HUVEC treated with NT siRNA. Images acquired at 5 min intervals over 240 min. Scale bar: 20μm.

**Supplementary Movie 4 (For Fig. 3K). SUN2 depletion disrupts coordinated migration of endothelial cells.** Movies acquired by phase-contrast confocal imaging of HUVEC treated with SUN2 siRNA. Images acquired at 5 min intervals over 240 min. Scale bar: 20μm.

**Supplementary Movie 5 (For Fig. 4E). Microtubule dynamics in control endothelial cells.** Movies acquired using spinning disc microscopy on EB3-mNeonGreen expressing HUVEC treated with NT siRNA. Images acquired at 470ms intervals over 60 sec. Black arrows indicate shrinking microtubules. Scale bar: 5μm.

**Supplementary Movie 6 (For Fig. 4E). SUN2 depletion increases microtubule stability in endothelial cells.** Movies acquired using spinning disc microscopy on EB3-mNeonGreen expressing HUVEC treated with SUN2 siRNA. Images acquired at 470ms intervals over 60 sec. Black arrows indicate shrinking microtubules. Scale bar: 5μm.

### SUPPLEMENTAL TABLES

#### **Supplementary Table S1:** Primers

| Primer | Application | Forward sequence (5'-3') | Reverse sequence (5'-3') |
|---|---|---|---|
| *Sun2 tm1a* | PCR (Mouse genotyping) | ACTATCCCGACCGCCTTACT | TAGCGGCTGATGTTGAACTG |
| *Sun2 floxed* | PCR (Mouse genotyping) | ATTGGGTGTCTCTACATCAAAGCTC | CCCCAGACAGAAGTGCCTAA |
| *FlpO* | PCR (Mouse genotyping) | TGAGCTTCGACATCGTGAAC | TCAGCATCTTCTTGCTGTGG |
| *Cdh5-CreERT2* | PCR (Mouse genotyping) | TCCTGATGGTGCCTATCCTC | CCTGTTTTGCACGTTCACCG |
| *sun2* | PCR (Zebrafish genotyping) | CTGAGAGCAGAGCTGTCTGATT | TGCTCCCTTTCTCATGTGCT |
| *MGP* | RT-qPCR (HUVEC) | CCTCAGCAGAGATGGAGAGCTA | ATGGCGTAGCGTTCGCAAAGTC |
| *LTBP1* | RT-qPCR (HUVEC) | TGAATGCCAGCACCGTCATCTC | CTGGCAAACACTCTTGTCCTCC |
| *ABI3BP* | RT-qPCR (HUVEC) | CCTTCTACACCTAAACGACGCC | GGTGTTGTCCATGTAGGTTCAGG |
| *FBLN5* | RT-qPCR (HUVEC) | CTGCTGGATGACAACCGAAGCT | GATAAGGCTCCTCACAGCGGAT |
| *ITGA9* | RT-qPCR (HUVEC) | TCAGGTCACAGAGAAGCTGCAG | CACATGCTCACTGAGGCTGTAG |
| *ITGB3* | RT-qPCR (HUVEC) | CATGGATTCCAGCAATGTCCTCC | TTGAGGCAGGTGGCATTGAAGG |
| *COL3A1* | RT-qPCR (HUVEC) | TGGTCTGCAAGGAATGCCTGGA | TCTTTCCCTGGGACACCATCAG |
| *FBN1* | RT-qPCR (HUVEC) | GGATACACAGGTGATGGCTTCAC | GTCGCATTCACAGCGGTATCCT |
| *UNC5B* | RT-qPCR (HUVEC) | GCTCGACTCTAAGAACTGCACAG | TGAGGATTGCCACGACCACGAA |
| *ADAMTSL2* | RT-qPCR (HUVEC) | CACATCCTGCAAGCTCACTGAC | CAGATGCCACACTTGTCCAGTG |
| *MMP9* | RT-qPCR (HUVEC) | GCCACTACTGTGCCTTTGAGTC | CCCTCAGAGAATCGCCAGTACT |
| *GAPDH* | RT-qPCR (HUVEC) | CAGCAAGAGCACAAGAGGAAGAGA | TTGATGGTACATGACAAGGTGCGG |

### Supplementary Table S2: Antibodies

| Primary antibodies | Application | Source | Catalog number |
|---|---|---|---|
| anti-VE-Cadherin | Mouse aorta staining | R&D Systems | AF1002 |
| anti-SUN2 | Mouse aorta staining | Abcam | ab124916 |
| anti-KLF4 | Mouse aorta staining, Western Blot | R&D Systems | AF3158 |
| anti-VE-Cadherin | HUVEC staining | Cell signaling | 2500S |
| anti-VE-Cadherin | HUVEC staining | Santa Cruz Biotechnology | sc-9989 |
| anti-GM130 | HUVEC staining | Abcam | ab52649 |
| anti-SUN2 | HUVEC staining | Sigma-Aldrich | MABT880 |
| anti-Paxillin | HUVEC staining, Western Blot | Abcam | ab32084 |
| anti-PECAM1 | HUVEC staining | Cell signaling | 3528S |
| anti-acetylated-α-tubulin | HUVEC staining | Sigma-Aldrich | T7451 |
| anti-α-tubulin | HUVEC staining | Cell signaling | 3873S |
| anti-Vinculin | Western Blot | Sigma-Aldrich | V9131 |
| anti-GAPDH | Western Blot | Cell signaling | 2118S |

| Secondary antibodies | Application | Source | Catalog number |
|---|---|---|---|
| Donkey anti-Rabbit IgG 488, 594, 647 | Immunofluorescence | Invitrogen | A21206, A21207, A31573 |
| Donkey anti-mouse IgG 488, 594, 647 | Immunofluorescence | Invitrogen | A21202, A21203, A31571 |
| Donkey anti-goat IgG 488, 594, 647 | Immunofluorescence | Invitrogen | A11055, A11058, A21447 |

| | | | |
|---|---|---|---|
| Donkey anti-goat IgG HRP | Western Blot | Invitrogen | PA1-28664 |
| Goat anti-rabbit IgG HRP | Western Blot | Invitrogen | 31460 |
| Goat anti-mouse IgG HRP | Western Blot | Invitrogen | 31430 |

**Supplementary Table S3:** siRNA and morpholinos

| siRNA/Morpholino | Application | Sequence (5'-3') | Source | Catalog number |
|---|---|---|---|---|
| NT morpholino | Zebrafish injection | CCTCTTACCTCAGTTACAATTTATA | GeneTools, LLC | N/A |
| *sun2* morpholino | Zebrafish injection | ATCTTGTGCTTCGTCTTGACATC | GeneTools, LLC | N/A |
| NT siRNA | HUVEC transfection | N/A | Dharmacon | D-001810-10-20 |
| *SUN2* siRNA | HUVEC transfection | N/A | Life technologies | 4392420, s24467 |
| *SYNE1* siRNA | HUVEC transfection | N/A | Dharmacon | M-014039-02-0005 |
